# Combination of AID2 and BromoTag expands the utility of degron-based protein knockdowns

**DOI:** 10.1101/2024.03.20.586026

**Authors:** Yuki Hatoyama, Moutushi Islam, Adam G. Bond, Ken-ichiro Hayashi, Alessio Ciulli, Masato T. Kanemaki

## Abstract

Acute protein knockdown is a powerful approach to dissecting protein function in dynamic cellular processes. We previously reported an improved auxin-inducible degron system, AID2, but recently noted that its ability to induce degradation of some essential replication factors, such as ORC1 and CDC6, was not enough to induce lethality. Here, we present combinational degron technologies to control two proteins and enhance target depletion. For this purpose, we initially compared PROTAC-based degrons, dTAG and BromoTag, with AID2 to reveal their key features and then demonstrated control of cohesin and condensin with AID2 and BromoTag, respectively. We developed a double-degron system with AID2 and BromoTag to enhance target depletion and accelerate depletion kinetics and demonstrated that both ORC1 and CDC6 are pivotal for MCM loading. Finally, we found that co-depletion of ORC1 and CDC6 by the double-degron system completely suppressed DNA replication, and the cells entered mitosis with single-chromatid chromosomes, indicating DNA replication was uncoupled from the cell cycle control. Our combinational degron technologies will expand the application scope for functional analyses.

## Introduction

Studies of protein function in living cells are greatly helped by conditional loss-of-function experiments. For this purpose, siRNA-based mRNA depletion and Cre-based conditional knockout have been employed for decades (Elbashir *et al*, 2001; Gu *et al*, 1993). However, target-protein depletion by these technologies is relatively slow because it depends on the protein’s half-life. For studying dynamic cellular processes such as the cell cycle, gene regulation and differentiation, rapid depletion of the target protein is crucial to capture the primary defect before the accumulation of secondary defects (Jaeger & Winter, 2021; Kanemaki, 2022). All eukaryotic cells are equipped with the ubiquitin–proteasome system (UPS), which degrades proteins within a few minutes to hours (Kleiger & Mayor, 2014). Therefore, protein knockdown by a conditional degron through the UPS provides an ideal methodology to study the rapid depletion of target proteins in living cells.

We previously developed one of the major conditional degron systems, auxin-inducible degron (AID), by transplanting a plant-specific degradation pathway into non-plant cells (Nishimura *et al*, 2009). For inducing target degradation, a degron tag derived from *Arabidopsis thaliana* IAA17 (e.g. mini-AID (mAID) and AID*) is fused to a protein of interest in cells expressing F-box protein *Oryza sativa* TIR1 (OsTIR1), which forms a SKP1–CUL1–F-box (SCF) E3 ligase with the endogenous components (Morawska & Ulrich, 2013; Natsume *et al*, 2016). A natural auxin, indole-3-acetic acid (IAA), added to the culture medium binds to OsTIR1 and, subsequently IAA-bound SCF–TIR1 recognizes and poly-ubiquitylates the degron for rapid degradation via the UPS. The original AID system showed a leaky degradation by SCF–OsTIR1 even without IAA addition, and the concentrations of IAA required for inducing target degradation were high (100 to 500 µM), which potentially exhibits toxicity in some cells. We and others recently overcame these problems by establishing an AID version 2 (AID2) system, in which we employed an OsTIR1(F74G/A) mutant and an auxin analogue, 5-Ph-IAA (**Figs 1A** and **S1A**) (Nishimura *et al*, 2020; Yesbolatova *et al*, 2020). We have successfully shown that AID2 allowed us to degrade proteins involved in DNA replication for phenotypic analyses (Klein *et al*, 2021; Lim *et al*, 2023; Liu *et al*, 2023; Saito *et al*, 2022). However, we recently noted that there were instances where target depletion by AID2 was not sufficient for phenotypic studies. In this study we sought to establish a methodology to enhance target depletion by AID2.

**Figure 1.**
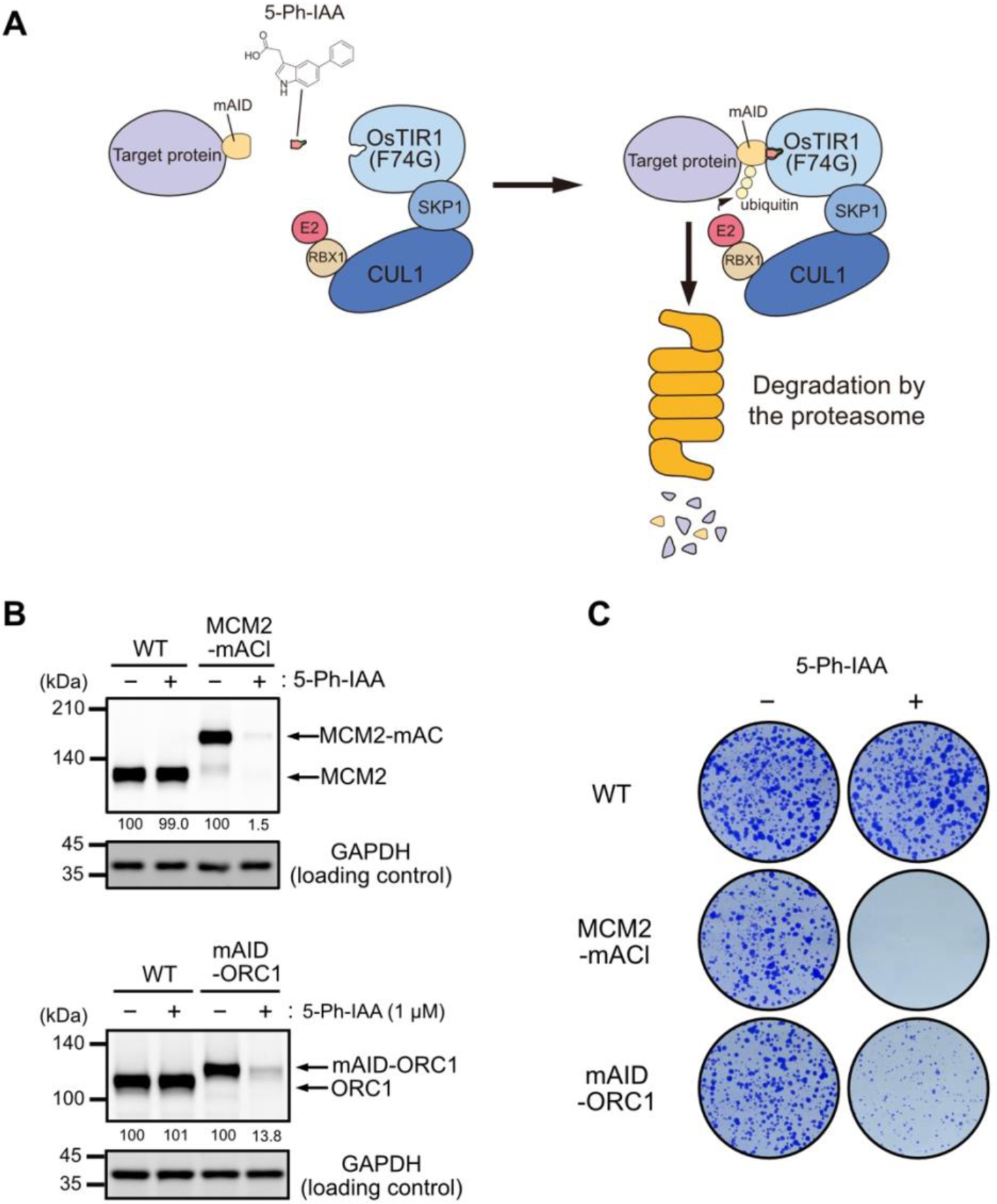
AID2-mediated depletion of MCM2 and ORC1 in HCT116. (**A**) Schematic illustration of the targeted protein degradation by the AID2 system. (**B**) Depletion of MCM2-mAID-mClover (MCM2-mAC) and mAID-ORC1. The parental HCT116 wild-type (WT), MCM2-mAC and mAID-ORC1 cells were treated with 1 µM 5-Ph-IAA for 6 h. Relative MCM2 and ORC1 levels taking the DMSO-treated control as 100% are shown under each blot. Each data point was normalized with the corresponding GAPDH loading control. Proteins were detected by anti-MCM2, -GAPDH and -tubulin antibodies. (**C**) Colony formation of the parental HCT116 WT, MCM2-mAC and mAID-ORC1 cells. The indicated cells were cultured in the presence or absence of 1 µM 5-Ph-IAA for 7 days. Colonies were stained with crystal violet.

Recently, other conditional degron technologies have been developed based on immunomodulatory drugs (IMiDs) or proteolysis-targeting chimeras (PROTACs) that utilise an endogenous E3 ligase such as CRL4–CRBN and CRL2–VHL (Bond *et al*, 2021; Bouguenina *et al*, 2023; Buckley *et al*, 2015; Koduri *et al*, 2019; Nabet *et al*, 2018; Nowak *et al*, 2021; Yamanaka *et al*, 2020). Because IMiDs inevitably induce off-target proteolysis of CRBN neo-substrates, PROTAC-based technologies have been an attractive choice for many researchers. Among PROTAC-based degrons, dTAG technology employs a PROTAC degrader, dTAG-13 or dTAGv-1, and FKBP12(F36V) as a degron (**Fig. S1A**) (Nabet *et al*, 2020; Nabet *et al*., 2018). A protein fused with FKBP12(F36V) (hereafter the degron tag is also called dTAG) is recruited to CRL4–CRBN or CRL2–VHL in the presence of dTAG-13 or dTAGv-1, respectively, for rapid degradation. BromoTag and a similar BRD4-based degron were recently reported (Bond *et al*., 2021; Nowak *et al*., 2021). BromoTag utilizes a PROTAC degrader AGB1 and an L387A mutant derived from the second BRD4 bromodomain as a degron (hereafter the degron tag is also called BromoTag) (**Fig. S1A**). A protein fused with BromoTag is recruited to CRL2–VHL in the presence of AGB1 for rapid proteolysis via the UPS. Both dTAG and BromoTag have been used in many studies, indicating that they are also promising conditional degrons (Evrin *et al*, 2023; Mylonas *et al*, 2021; Olsen *et al*, 2022; Weintraub *et al*, 2017). However, few studies have compared conditional degrons to discern their similarities and differences, and BromoTag has not yet been included for comparison (Bondeson *et al*, 2022; Noviello *et al*, 2023). Furthermore, the target proteins fused with each degron in these studies were overexpressed to varying levels, impacting degradation kinetics and complicating the comparison. Ideally, a comparison of different degron systems should be conducted with a cell line expressing the same level of target protein.

To achieve the goal of enhancing target depletion by AID2, we proposed combining AID2 with another degron system. We initially compared AID2, dTAG and BromoTag using a single GFP reporter containing three degron tags in tandem to understand their similarities and differences. We found that all degron systems achieve rapid reporter depletion. However, AID2 exhibits superior performance in terms of depletion efficiency, kinetics and reversible expression recovery in HCT116 and hTERT-RPE1 cells. Subsequently, we showed that two proteins can be independently and simultaneously depleted using AID2 and BromoTag. Finally, we showed that a double-degron system with mAID-BromoTag enhances target depletion, accelerates depletion kinetics, and confers strong phenotypic defects. By using this double-degron system, we succeed in showing that both ORC1 and CDC6 are pivotal for the MCM-loading prerequisite for DNA replication. Furthermore, we achieved complete suppression of DNA replication by co-depleting ORC1 and CDC6, leading cells to enter mitosis without DNA replication.

## Results

### AID2-mediated ORC1 depletion does not result in a strong growth defect

We previously established an improved version of the auxin-inducible degron (AID) system, namely AID2, and reported that degron-fused proteins were sharply induced for rapid degradation after the addition of 5-Ph-IAA (**Fig. 1A**) (Yesbolatova *et al*., 2020). Because our group is interested in DNA replication, we applied AID2 to control the expression of essential replication factors in human colorectal cancer HCT116 cells by biallelically tagging the endogenous target genes using CRISPR-Cas9 (Lim *et al*., 2023; Liu *et al*., 2023; Saito *et al*., 2022). When we applied AID2 to the MCM2 subunit of the replicative MCM2–7 helicase, we successfully depleted MCM2 C-terminally fused with mAID-mClover (MCM2-mACl) upon addition of 5-Ph-IAA (**Fig. 1B, upper blots**). Consequently, MCM2-mACl cells treated with 5-Ph-IAA stopped growing and did not form any colonies, suggesting that they did not carry out DNA replication (**Fig. 1C**).

ORC1 is a subunit of the ORC1–6 complex, which plays a pivotal role in loading MCM2–7 to chromosomal DNA, and loss of ORC1 causes lethality in budding yeast (Klemm & Bell, 2001). However, there are conflicting reports on whether ORC1 is essential for DNA replication or not in human cells (Chou *et al*, 2021; Shibata *et al*, 2016). To clarify this issue, we generated ORC1-degron cells with AID2 by fusing mAID to the N-terminus of ORC1. The mAID-ORC1 protein was efficiently depleted upon addition of 5-Ph-IAA (**Fig. 1B, lower blots**). Unexpectedly, mAID-ORC1 cells grew slowly and formed small colonies, suggesting that ORC1-depleted cells carried out DNA replication even though the cells were defective (**Fig. 1C**). We interpreted this result as suggesting that ORC1 was likely to be required for human DNA replication, but the depletion by AID2 was not enough to cause lethality, and the required amount of ORC1 for DNA replication was very low compared with that of MCM2. We found a similar case with another replication factor, CDC6 (described later). These findings motivated us to enhance depletion by the AID2 system.

### Understanding similarities and differences of AID2, dTAG and BromoTag

To overcome the challenges mentioned above, we sought to combine AID2 and another degron system. We became interested in the recently reported PROTAC-based degrons, dTAG and BromoTag (**Fig. S1A**) (Bond *et al*., 2021; Nabet *et al*., 2020; Nabet *et al*., 2018; Nowak *et al*., 2021). Even though AID2, dTAG and BromoTag have been used in cell biology, there are only a few studies comparing these systems (Bondeson *et al*., 2022; Noviello *et al*., 2023). To understand the similarities and differences among AID2, dTAG and BromoTag systems, we constructed a GFP reporter containing the three degrons in tandem (**Fig. 2A**). These degrons (dTAG, BromoTag and mAID, respectively) were connected with a long linker (11–12 amino-acid chain composed of G, A and S) to ensure that they can each be readily accessed by the corresponding E3 ubiquitin ligase enzyme complex. In order to focus on nuclear proteins such as ORC1, we also fused a nuclear localization signal (NLS) to the C-terminus and confirmed its nuclear localization (**Fig. S2A**). We transfected a transposon plasmid encoding the reporter gene to parental HCT116 wild-type (WT) cells expressing OsTIR1(F74G) from the safe harbour *AAVS1* locus and selected stable cells as previously reported (**Fig. 2A**) (Yesbolatova *et al*., 2020).

**Figure 2.**
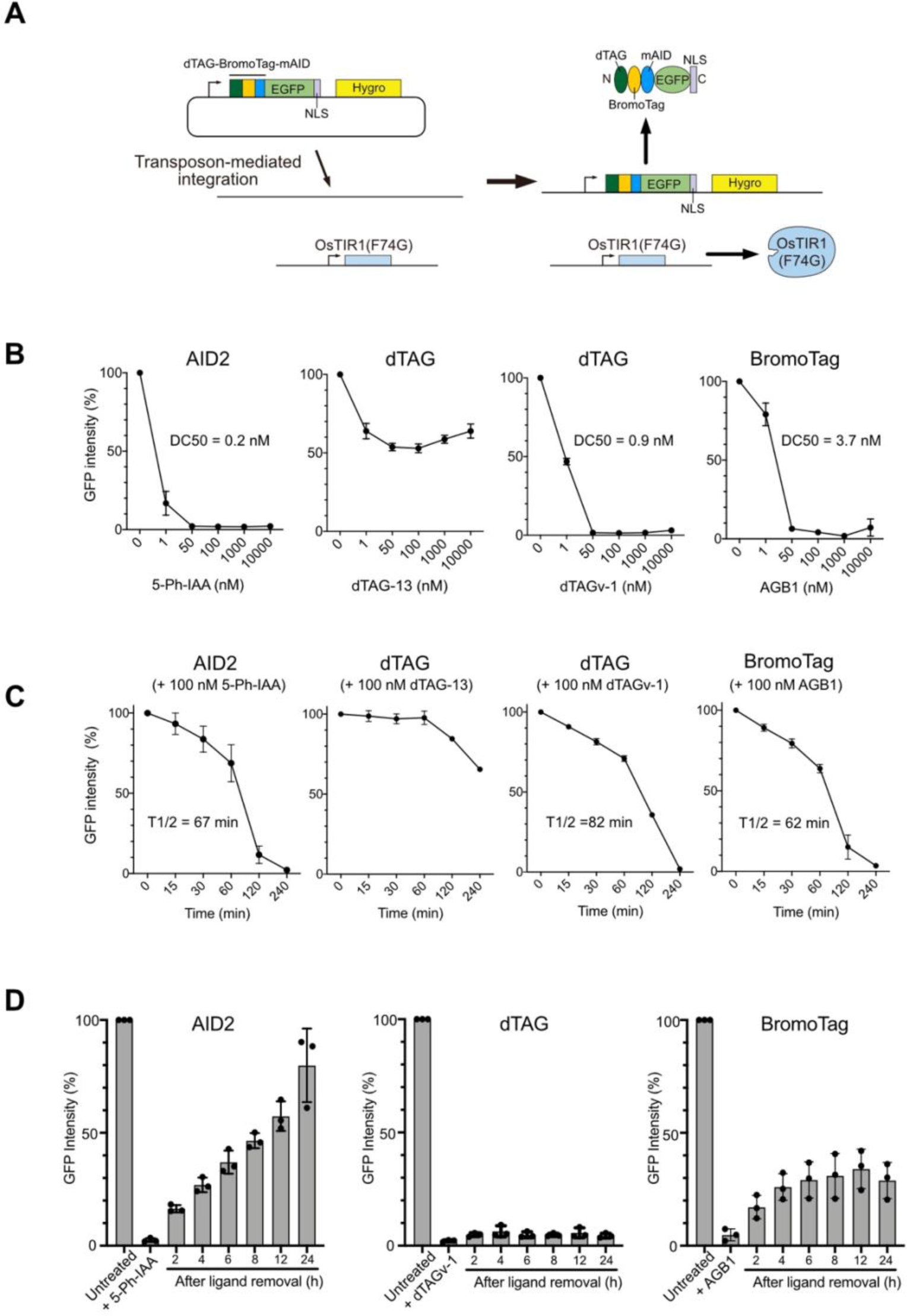
Comparing the AID2, dTAG and BromoTag systems using the reporter HCT116 cells. (**A**) Schematic illustration showing the strategy to generate the reporter HCT116 cell line expressing OsTIR1(F74G) and the GFP reporter. (**B**) Dose–response of reporter depletion. The reporter cells were treated with the indicated concentrations of each ligand for 4 h. GFP intensity was analysed taking the mock-treated cells as 100% (mean ± SD, n = 4). The DC50 values were calculated with the non-linear regression model on GraphPad Prism 8. (**C**) Time-course depletion of the reporter. The reporter cells were treated with 100 nM of the indicated ligand. Samples were taken at the indicated time points, and the GFP intensity was analysed taking the mock-treated cells as 100% (mean ± SD, n = 4). The T1/2 was calculated with the non-linear regression model on GraphPad Prism 8. (**D**) Re-expression of the reporter after depletion by the AID2, dTAG or BromoTag system. The reporter cells were treated with 100 nM 5-Ph-IAA, dTAGv-1 or AGB1 for 4 h before medium change. Samples were taken at the indicated time points, and the GFP intensity was analysed taking the mock-treated cells as 100% (mean ± SD, n = 3).

We initially tested ligand concentrations required for inducing reporter degradation (**Fig. 2B**). The reporter cells were treated with a differential concentration of inducing ligand for 4 h, and subsequently, the GFP reporter level was monitored using a flow cytometer. In the case of AID2, efficient depletion was achieved with the lowest concentration (DC50 = 0.2 nM 5-Ph-IAA). There are two inducing ligands for the dTAG system, dTAG-13 and dTAGv-1, which recruit CRL4–CRBN and CRL2–VHL E3 ligase, respectively (**Fig. S1A**) (Nabet *et al*., 2020; Nabet *et al*., 2018). We found that dTAG-13 did not efficiently deplete the GFP reporter in HCT116 cells. On the other hand, dTAGv-1 worked as expected and achieved almost complete depletion at 50 nM (DC50 = 0.9 nM). The reason why dTAG-13 did not work efficiently in HCT116 cells might be because the expression levels of CRL4–CRBN, which recognizes dTAG in the presence of dTAG-13, were lower than those of the other E3 ubiquitin ligases required by 5-Ph-IAA, dTAGv-1 and AGB1 (**Fig. S1B**) (Bekker-Jensen *et al*, 2017). AGB1 for BromoTag also induced depletion of the GFP reporter, albeit slightly less efficiently (DC50 = 3.7 nM). To study whether the trends we found were also true in another cell line, we generated a similar reporter cell line in a normal human cell line, hTERT-RPE1, and found a similar trend, although dTAG-13 was more effective in this cell line (**Fig. S2B**). Unlike the AID2 system, PROTAC dTAGs and AGB1 showed the hook effect at high doses of ligands, a characteristic feature of heterobifunctional degraders **(Figs 2B and S2B).**

Next, we investigated depletion kinetics by treating the HCT116 reporter cells with 100 nM ligand and then took time-course samples (**Fig. 2C**). We found that T1/2 was 67, 82 and 62 min when treated with 5Ph-IAA, dTAGv-1 and AGB1, respectively, indicating AID2 and BromoTag comparably induced slightly faster depletion than dTAG with dTAGv-1. We also studied the depletion kinetics in the hTERT-RPE1 reporter cells and found T1/2 of AID2, dTAG (with dTAGv-1) and BromoTag were 13, 62 and 42 min, respectively (**Fig. S2C**). These results indicated that AID2 and BromoTag depleted the GFP reporter quicker than dTAG in both HCT116 and hTERT-RPE1 cells.

An ideal inducible degron system should operate reversibly. To investigate whether the GFP reporter can be re-expressed after depletion, we initially treated the HCT116 GFP reporter cells with 100 nM 5-Ph-IAA, dTAGv-1 or AGB1 for 4 h to induce GFP reporter depletion. Subsequently, the cells were washed and incubated in fresh medium without each ligand to monitor re-expression of the GFP reporter (**Fig. 2D**). As previously reported, the cells treated with 5-Ph-IAA recovered GFP reporter expression over time, with 8 h required to achieve 50% recovery (**Fig. 2D, AID2**) (Yesbolatova *et al*., 2020). Unexpectedly, the cells treated with dTAGv-1 did not show any recovery until 24 h (**Fig. 2D, dTAG**). The cells treated with AGB1 recovered up to about 30% at 6 h, but no further recovery was observed (**Fig. 2D, BromoTag**). We found similar results in expression recovery in hTERT-RPE1 cells (**Fig. S2D**). These results indicate that PROTAC-based degrons exhibited poor recovery after target depletion, suggesting that dTAGv-1 and AGB1 continued to bind CRL2–VHL E3 ligase even after medium exchange, maintaining target degradation, consistent with their catalytic substoichiometric mode of action at low concentrations. Considering the results of ligand concentration (**Figs 2B** and **S2B**), depletion kinetics (**Figs 2C** and **S2C**) and expression recovery (**Figs 2D** and **S2D**), we decided to combine AID2 and BromoTag in the following experiments.

### Independent and simultaneous depletion of two proteins

Because we found that AID2 and BromoTag achieved rapid depletion of the GFP reporter (**Fig. 2C**), we asked whether two proteins can be independently depleted with AID2 and BromoTag. For this purpose, we chose two structural maintenance-of-chromosome (SMC) complexes, cohesin and condensin, which are involved in sister-chromatid cohesion and mitotic chromosome formation, respectively (Jeppsson *et al*, 2014). We fused mAID-mClover to the C-terminus of the endogenous RAD21 subunit of cohesin as previously reported (**Fig. S3A**) (Natsume *et al*., 2016). Similarly, we introduced BromoTag-mCherry2 to the endogenous SMC2 subunit of condensin. The established HCT116 cell line expressed both RAD21-mAID-mClover (RAD21-mACl) and SMC2-BromoTag-mCherry2 (SMC2-BTCh), which are induced for rapid degradation in the presence of 5-Ph-IAA and AGB1, respectively (**Fig. 3A**). Initially, we confirmed the expression of RAD21-mACl and SMC2-BTCh by fluorescence microscopy (**Fig. S3B, control**).

**Figure 3.**
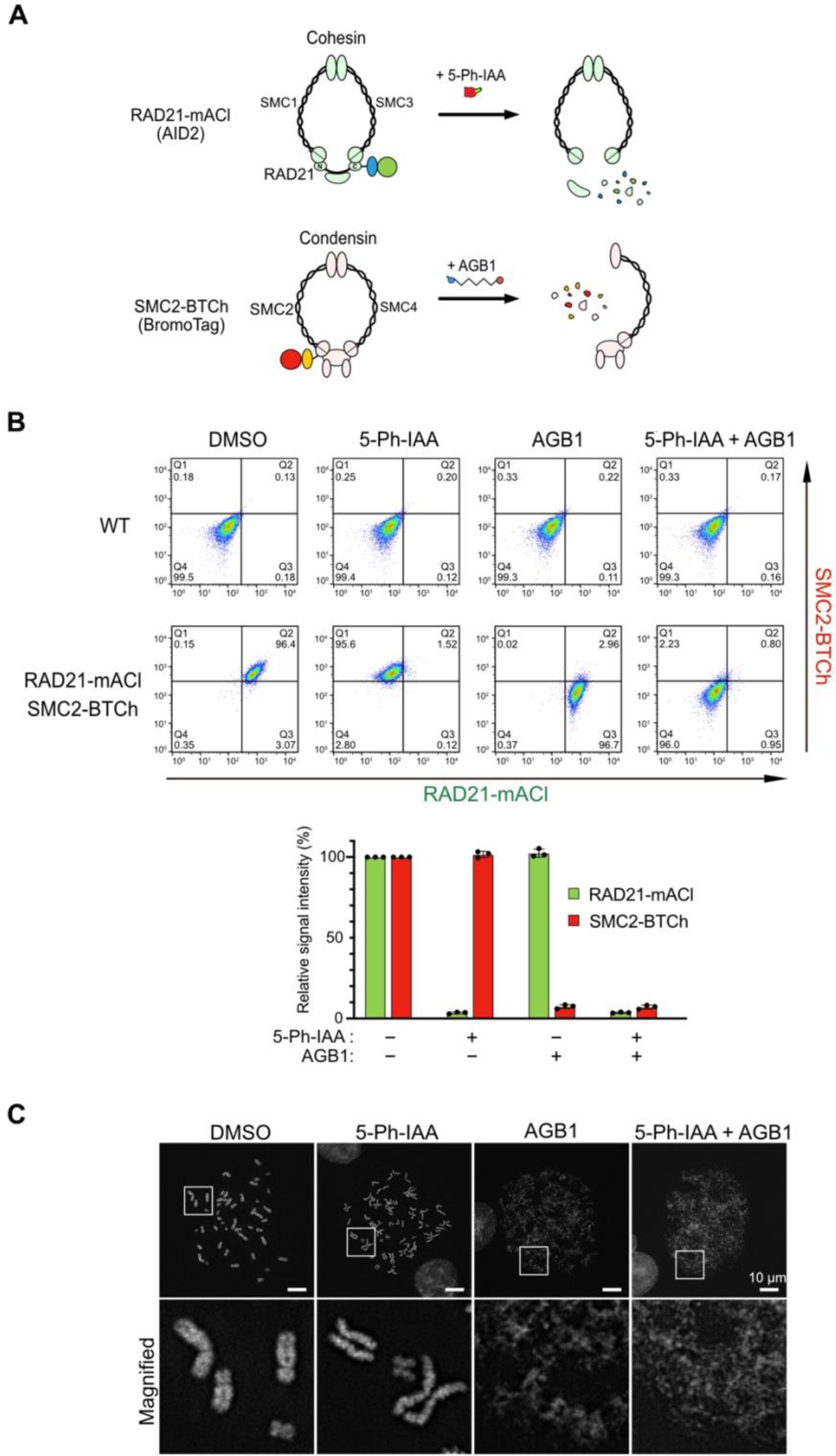
Independent and simultaneous depletion of RAD21-mAID-mClover (RAD21-mACl) and SMC2-BromoTag-mCherry2 (SMC2-BTCh). (**A**) Schematic illustration showing degradation of RAD21-mACl and SMC2-BTCh in the cohesin and condensin complexes, respectively. (**B**) Top, density plots of the cells treated with 1 µM 5-Ph-IAA, 0.5 µM AGB1 and both for 2 h. The X and the Y axes are the signal intensity of RAD21-mACl and SMC2-BTCh, respectively. Quadrant gates (Q1–Q4) were set manually to include 99.5% of the single cells into Q4 area using a negative control (DMSO-treated WT cells). Bottom, the density plot data were quantified to make this graph (mean ± SD, n=3). The signal intensities in each condition were normalized with DMSO-treated control cells. (**C**) Representative DAPI-stained chromosomes from RAD21-mACl/SMC2-BTCh cells after treating them with the indicated ligand for 3 h. The scale bars indicate 10 µm.

Before depleting RAD21-mACl and SMC2-BTCh in this cell line, we optimized inducing concentrations of 5-Ph-IAA and AGB1. We cultured the parental HCT116 WT cells in the presence of 5-Ph-IAA or AGB1 for 7 days (**Fig. S3C**). We found that 1 µM 5-Ph-IAA did not affect cellular proliferation and colony formation as previously reported (Yesbolatova *et al*., 2020). On the other hand, the cells grown in 1 µM AGB1 formed smaller colonies, suggesting that AGB1 affected cellular proliferation. The cells grown in 0.5 µM AGB1 formed colonies as in the control DMSO-added culture. Therefore, we decided to employ 1 µM 5-Ph-IAA and 0.5 µM AGB1 for the following experiments.

We treated the cells with 5-Ph-IAA or AGB1 independently, or together. After treating the cells, the expression level of RAD21-mACl and SMC2-BTCh was monitored by flow cytometry and microscopy (**Figs 3B** and **S3B**). When treated with 5-Ph-IAA or AGB1, each target protein was depleted without affecting the other protein. The addition of both 5-Ph-IAA and AGB1 combined resulted in depletion of both RAD21-mACl and SMC2-BTCh. These data clearly show that two endogenous proteins were independently or simultaneously depleted with the orthogonal AID2 and BromoTag systems.

Subsequently, we investigated phenotypic defects in chromosomal structures after the depletion of RAD21-mACl and SMC2-BTCh. For this purpose, we looked at mitotic chromosomes after RAD21-mACl and SMC2-BTCh depletion (**Fig. 3C**). Chromosomes in the 5-Ph-IAA-treated cells (RAD21-depleted cells) showed loss of sister-chromatid cohesion (**Fig. 3C, 5-Ph-IAA**) (Sonoda *et al*, 2001; Yesbolatova *et al*., 2020). Chromosomes in AGB1-treated cells (SMC2-depleted cells) showed the rod-like structure of mitotic chromosomes was highly disorganized, as previously reported for loss of condensin (**Fig. 3C, AGB1**) (Boteva *et al*, 2020; Takagi *et al*, 2018). Furthermore, chromosomes in the 5-Ph-IAA- and AGB1-treated cells (RAD21- and SMC2-depleted cells) were even more disorganized than those treated with AGB1 alone, possibly because of an additive effect of cohesion and structural losses (**Fig. 3C, 5-Ph-IAA + AGB1**). These results demonstrate that two proteins can be independently and simultaneously depleted in a single cell by utilizing AID2 and BromoTag, allowing dissection of the functional consequences of knocking down two proteins individually or concurrently.

### Enhancing target protein depletion by an AID2-BromoTag double-degron system

As we presented in **Fig. 1**, there are cases in which target depletion by AID2 does not result in a strong phenotypic defect. We hypothesized that a double-degron tag composed of mAID and BromoTag in tandem would enhance target protein depletion, thus causing stronger phenotypic defects. To test this idea, we investigated whether the GFP reporter used in **Fig. 2** can be depleted better by treating the reporter cells with both 5-Ph-IAA and AGB1. We initially identified that treatment with 0.2 nM 5-Ph-IAA or 3.8 nM AGB1 caused about 50% reduction in expression of the GFP reporter (48.3% ± 3.3 or 50% ± 1.5, respectively) (**Fig. 4A, bars 2 and 3**). Subsequently, we treated the GFP reporter cells with both 0.2 nM 5-Ph-IAA and 3.8 nM AGB1 and found that the GFP expression level was 24.7% ± 2.6, indicating an additive degradation by AID2 and BromoTag (**Fig. 4A, bar 4**). We were encouraged by the additive effect in the expression level and asked whether depletion kinetics can also be accelerated. To answer this question, we treated the GFP reporter cells with 1 µM 5-Ph-IAA, or 0.5 µM AGB, or both, and then took time-course samples. **Fig. 4B** clearly shows that depletion kinetics was accelerated by treating both ligands (T1/2 = 25.7 min). Single ligand treatment achieved comparable depletion kinetics (5-Ph-IAA T1/2 = 66.4 min, AGB1 T1/2 = 53.6 min; not statistically significant by paired t test). These results demonstrated that the AID2-BromoTag double-degron system induced enhanced degradation and accelerated depletion kinetics compared with the case of the single degron.

**Figure 4.**
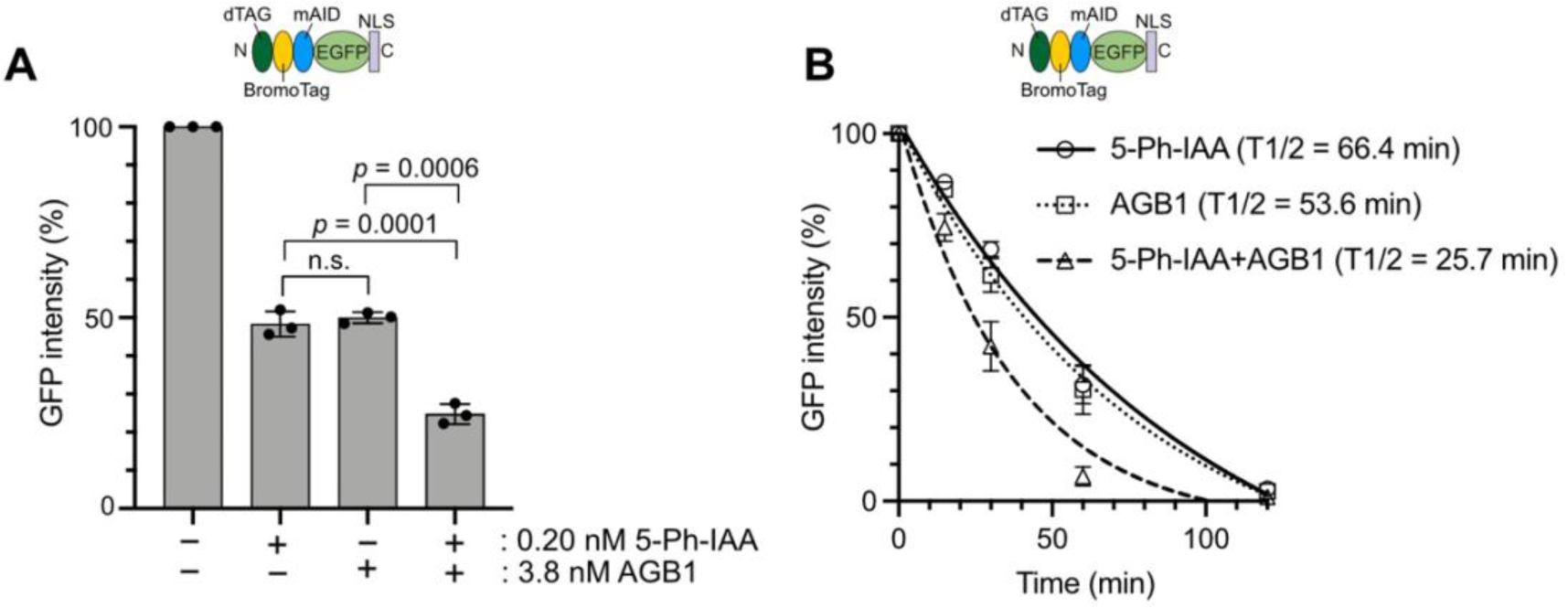
Enhancing reporter depletion by a double-degron system with AID2 and BromoTag. (**A**) The reporter cells used in Figs 2 and 3 were treated with 0.2 nM 5-Ph-IAA and/or 3.8 nM AGB1 for 4 h. GFP intensity was monitored taking the mock-treated cells as 100% (mean ± SD, n = 3). (**B**) Depletion kinetics of the reporter depletion in cells treated with 1 µM 5-Ph-IAA, 0.5 µM AGB1 or both. The GFP intensity was monitored at 0, 15, 30, 60, 120 and 240 min taking the mock-treated cells as 100% (mean± SD, n = 3). The data were fitted with one-phase decay.

### ORC1 and CDC6 depletion by the AID2-BromoTag double-degron system

Considering the results with the GFP reporter shown above, we then investigated whether AID2-BromoTag will give better depletion and phenotypic defects for ORC1. We fused mAID-BromoTag to the N-terminus of endogenous ORC1(mAB-ORC1). The mAB-ORC1 cells were treated with 5-Ph-IAA, AGB1 or both for 2 h (**Fig. 5A**). Compared with the mAB-ORC1 expression level treated with 5-Ph-IAA or AGB1 (16.6% or 7.7%, respectively), expression was further reduced when treated with both (3.9%), similar to the case observed with the GFP reporter in **Fig. 4**. Subsequently, we investigated the depletion kinetics of mAB-ORC1 treated with 5-Ph-IAA, AGB1 or both, and then took time-course samples (**Fig. 5B, C**). At every time point, the cells treated with both ligands showed better depletion of mAB-ORC1, indicating that mAB-ORC1 depletion was enhanced and accelerated by treating with 5-Ph-IAA and AGB1 together.

**Figure 5.**
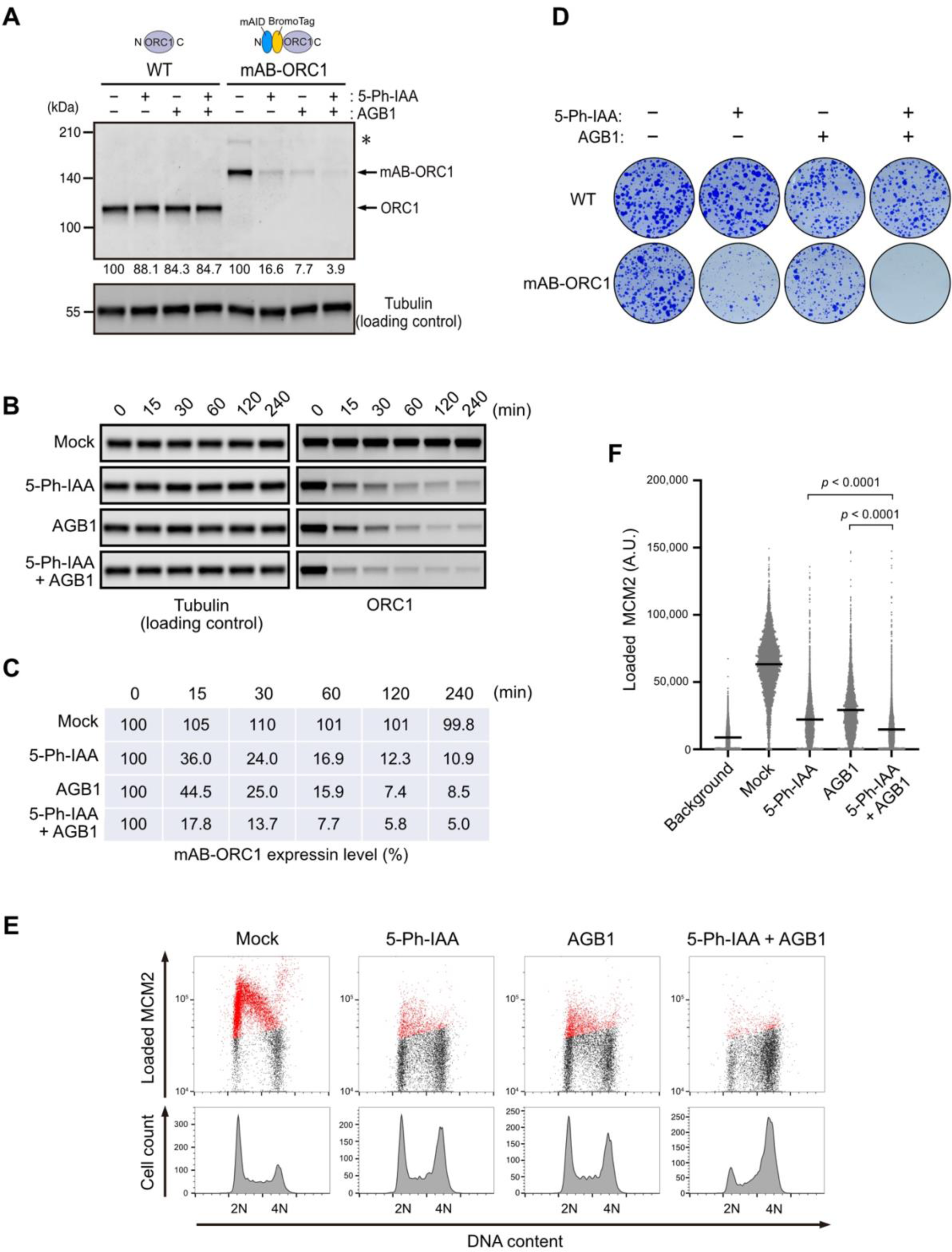
Double-degron with mAID and BromoTag enhances ORC1 depletion and confers profound defects in DNA replication. (**A**) The parental HCT116 wild-type (WT) and mAID-BromoTag-ORC1 (mAB-ORC1) cells were treated with 1 µM 5-Ph-IAA, 0.5 µM AGB1 or both for 2 h. Proteins were detected by anti-ORC1 and -tubulin antibodies. Relative ORC1 levels taking the DMSO-treated control as 100% are shown under each blot. Each data point was normalized with the corresponding tubulin loading control. The asterisk indicates the HygR-P2A-mAID-BromoTag-ORC1 protein before self-cleavage at the P2A site. (**B**) Depletion kinetics of mAB-ORC1 in cells treated with 1 µM 5-Ph-IAA and/or 0.5 µM AGB1. Samples were taken at the indicated time points. (**C**) The blot data in panel B were quantified taking the 0 min sample as 100%. Each data point was normalized with the corresponding tubulin loading control. (**D**) Colony formation of the parental HCT116 WT and mAB-ORC1 cells. Cells were cultured in the presence or absence of 1 µM 5-Ph-IAA and/or 0.5 µM AGB1 for 7 days. Colonies were stained with crystal violet. (**E**) (Upper panels) Levels of chromatin-loaded MCM2 and DNA in mAB-ORC1 treated with 1 µM 5-Ph-IAA and/or 0.5 µM AGB1 for 24 h. The MCM2-positive cells are shown in red. (Lower panels) Cell count histogram for the same samples. (**F**) Levels of chromatin-loaded MCM2 in mAB-ORC1 cells treated with the indicated ligand (bars: mean, n > 5,000 cells). Cells were synchronized in M phase with 50 ng/mL nocodazole for 14 h and released into fresh media containing ligand. Cells were treated with 1 µM 5-Ph-IAA and/or 0.5 µM AGB1 2 h prior to nocodazole release. Samples were taken at 4 h after release when cells were in G1. Fixed cells stained without MCM2 antibody serve as the background. Statistical analysis was performed with a Kruskal–Wallis test.

Next, we studied the viability of the mAB-ORC1 cells by culturing them in the presence of 5-Ph-IAA, AGB1 or both (**Fig. 5D**). Cells grown with 5-Ph-IAA made small colonies as observed with mAID-ORC1 cells in **Fig. 1C**. Similarly, cells grown with AGB1 formed small colonies, although the colonies were slightly bigger than those of the 5-Ph-IAA-treated cells. In contrast, the cells with both 5-Ph-IAA and AGB1 did not form colonies, showing that mAB-ORC1 depletion caused lethality. ORC1 plays an essential role in loading MCM2–7 to chromosomal DNA in late M to G1 phases in budding yeast (Klemm & Bell, 2001). We investigated whether mAB-ORC1 depletion would affect MCM2–7 loading in human cells by looking at chromatin-bound MCM2 (**Fig. 5E**) (Haland *et al*, 2015). The mAB-ORC1 cells were cultured with 5-Ph-IAA, AGB1 or both for 24 h. Subsequently, chromatin-bound MCM2 was stained after extraction of chromatin-unbound MCM2. The level of chromatin-bound MCM2 shown in red was reduced in the cells treated with 5-Ph-IAA or AGB1 (**Fig. 5E**). However, chromatin-bound MCM2 was even more reduced in the cells treated with both 5-Ph-IAA and AGB1. We confirmed this notion by synchronizing cells and quantified the levels of chromatin-bound MCM2 in the G1 phase (**Fig. 5F**). Furthermore, the proportion of cells arrested in late S to G2 phases was the highest with 5-Ph-IAA and AGB1, suggesting these cells were experiencing strong defects in DNA replication with small amounts of chromatin-loaded MCM2–7 (**Fig. 5E**). Consistent with this interpretation, we observed 53BP1 foci showing that DNA damage was highly accumulated in the cells treated with both 5-Ph-IAA and AGB1 for 43 h, as compared with those treated with either 5-Ph-IAA or AGB1 (**Fig. S4A, B**). Considering these results, we concluded that ORC1 is essential for loading MCM2–7 onto chromatin DNA and thus is pivotal for DNA replication in human cells. Next, we wished to test the AID2-BromoTag double-degron system with another DNA replication factor. It has been reported that the original AID system was employed for depleting CDC6, which collaborates with ORC1–6 for loading MCM2–7 (Lemmens *et al*, 2018). However, the depletion was insufficient to observe defects in DNA replication. Therefore, we fused mAID-BromoTag to the N-terminus of the endogenous CDC6 protein (mAB-CDC6) and carried out similar experiments to those done for mAB-ORC1 (**Fig. S5**). When we treated mAB-CDC6 cells with 5-Ph-IAA, AGB1 or both for 2 h, we observed an additive depletion resulting in reduced expression of mAB-CDC6 down to 9% (**Fig. S5A**). Depletion of mAB-CDC6 was accelerated by treating with both 5-Ph-IAA and AGB1 (**Fig. S5B, C**). Importantly, treating with both 5-Ph-IAA and AGB1 was lethal to the mAB-CDC6 cells (**Fig. S5D**). In contrast, a single treatment with 5-Ph-IAA or AGB1 led to slow growth. Furthermore, treating with both 5-Ph-IAA and AGB1 induced strong defects in the loading of MCM2–7, arresting cells in late S to G2 phases with 53BP1 foci accumulation (**Figs S4** and **S5E, F**). Taken all together, we concluded that the double-degron system with AID2 and BromoTag is more effective than the respective single-degron at enhancing target depletion and inducing pronounced phenotypic defects. We also concluded that both ORC1 and CDC6 are required for the MCM loading, the essential process prerequisite for DNA replication.

### Tandem mAID tags are unstable even though they induce strong phenotypic defects

Because the double-degron system with AID2 and BromoTag enhanced target depletion, we wondered whether utilizing a tandem mAID tag would also enhance target depletion, as previously reported in yeast (Kubota *et al*, 2013; Nishimura & Kanemaki, 2014). We fused one, two or three copies of mAID (mAID, 2mAID, or 3mAID, respectively) to the N-terminus of ORC1 and compared them with each other (**Fig. 6**). We found that the expression level of 2mAID- and 3mAID-ORC1 was significantly reduced compared with that of mAID-ORC1 (**Fig. 6A, – 5-Ph-IAA**). This instability was also observed in cells expressing 2mAID-CDC6 (**Fig. S6A**). The reduced expression of 2mAID-ORC1 is due to instability of the 2mAID tag and is independent of leaky degradation by OsTIR1(F74G) (**Fig. S6B**). Even though the expression of 2mAID- and 3mAID-ORC1 was reduced, the cells proliferated normally in the absence of 5-Ph-IAA (**Fig. 6B, + DMSO**). Upon the addition of 5-Ph-IAA, 2mAID- and 3mAID-ORC1 were effectively depleted (**Fig. 6A**) and showed a lethal phenotype (**Fig. 6B, + 5-Ph-IAA**). As expected, the treated cells were defective in loading MCM2–7 onto chromatin and arrested in late-S to G2 phase (**Fig. 6C, D**). Therefore, we concluded that 2mAID and 3mAID allow for stronger phenotypic defects, with the caveat that they induce instability on the fusion protein.

**Figure 6.**
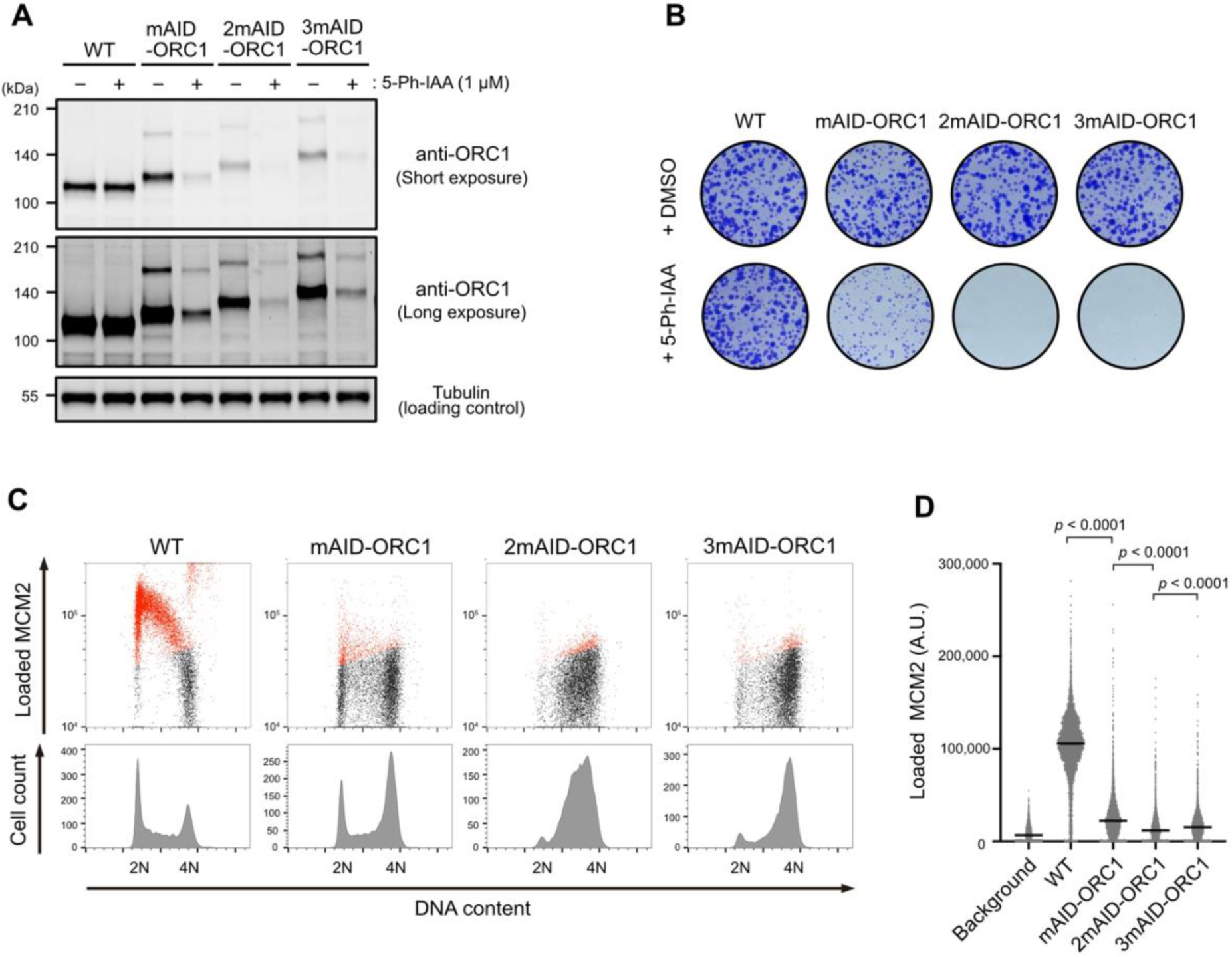
ORC1 depletion and phenotypic defects of mAID-, 2mAID- and 3mAID-ORC1 cells. (**A**) ORC1 level in cells treated with 1 µM 5-Ph-IAA for 2 h. (**B**) Colony formation of the indicated cells. Cells were cultured in the presence or absence of 1 µM 5-Ph-IAA for 7 days. Colonies were stained with crystal violet. (**C**) (Upper panels) Level of chromatin-loaded MCM2 and DNA in the indicated cells. Cells were cultured with 1 µM 5-Ph-IAA for 24 h. MCM2-positive cells are shown in red. (Lower panels) Cell count histogram for the same samples. (**D**) Levels of chromatin-loaded MCM2 in indicated cell lines treated with 5-Ph-IAA (bars: mean, n > 3,500 cells). Cells were synchronized in M phase with 50 ng/mL nocodazole for 14 h and released into fresh media containing 5-Ph-IAA. Cells were treated with 1 µM 5-Ph-IAA 2 h prior to nocodazole release. Samples were taken at 4 h after release when cells were in G1. Fixed WT cells stained without MCM2 antibody serve as the background. Statistical analysis was performed with a Kruskal–Wallis test.

### Complete inhibition of DNA replication leads to uncoupling DNA replication from the cell cycle control

Up until this point, we had succeeded in depleting ORC1 and CDC6 using the AID2-BromoTag double-degron system, and demonstrated their importance for MCM2–7 loading and cellular proliferation (**Figs 5** and **S5**). However, mAB-ORC1 and mAB-CDC6 cells replicated genomic DNA mostly and arrested around 4N (**Figs 5E** and **S5E**). The same was true when we used 2mAID-ORC1 (**Fig. 6C**). These results suggest two intriguing implications. First, human HCT116 cells do not have the licensing checkpoint, monitoring the amount of chromatin-bound MCM2–7 in the G1 phase because they carried out DNA replication with reduced chromatin-bound MCM2–7, consistent with our previous report (Saito *et al*., 2022). Second, the results suggest the ORC1 depletion was still not enough to completely suppress MCM loading and DNA replication. To clarify the second issue, we generated a double-mutant cell line for ORC1 and CDC6: mAB-ORC1 mAB-CDC6. We found that MCM loading was completely suppressed, arresting the cells at 2N when ORC1 and CDC6 were simultaneously depleted (**Figs 7A** and **S7A**). Furthermore, we observed that DNA replication was completely suppressed in a synchronous culture released from nocodazole arrest (**Fig. 7B**). These results indicate that ORC1 single depletion in **Figs 5** and **6** was not enough to suppress DNA replication completely, and leaky DNA replication still occurred.

**Figure 7.**
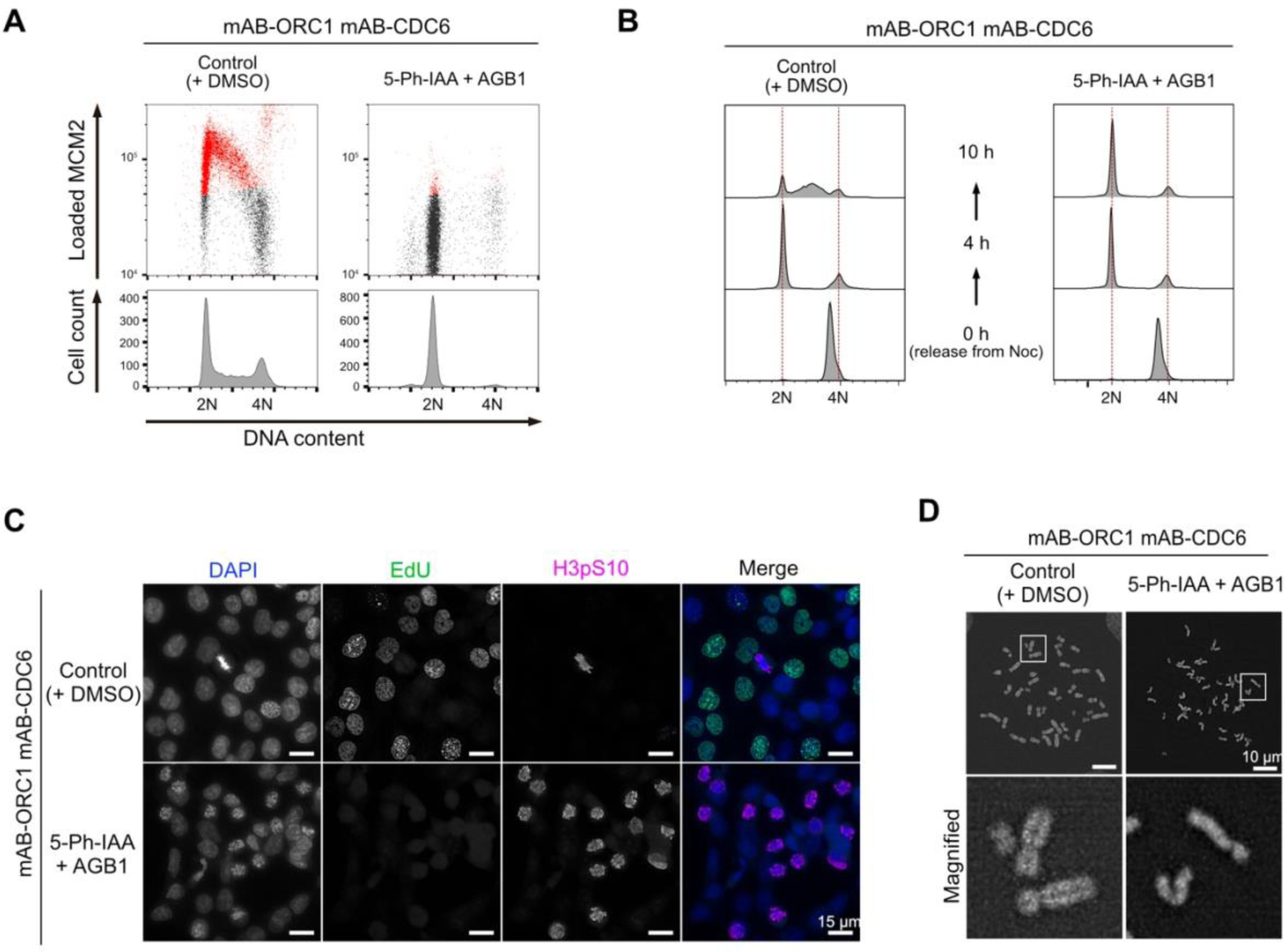
Complete inhibition of DNA replication in the mAB-ORC1 mAB-CDC6 cells leads to premature mitosis. (**A**) (Upper panels) Levels of chromatin-loaded MCM2 and DNA in the mAB-ORC1 mAB-CDC6 cells treated with 1 µM 5-Ph-IAA for 24 h. The MCM2-positive cells are shown in red. (Lower panels) Cell count histogram for the same samples. (**B**) Cell cycle progression of the mAB-ORC1 mAB-CDC6 cells. Cells were synchronized with 50 ng/mL nocodazole for 14 h and then released into fresh media containing 1 µM 5-Ph-IAA and 0.5 µM AGB1. Cells were treated with 5-Ph-IAA and AGB1 2 h prior to nocodazole release. Samples were taken at the indicated time points. (**C**) Fluorescent microscopic images of mAB-ORC1 mAB-CDC6 cells cultured with or without 1 µM 5-Ph-IAA and 0.5 µM AGB1 for 24 h. Cells were treated with 10 µM EdU for 30 min before fixation. Cells under mitosis were stained with anti-histone H3 p-Ser10 antibody. The scale bars indicate 15 µm. (**D**) DAPI-stained spread chromosomes after treating the mAB-ORC1 mAB-CDC6 cells with or without 1 µM 5-Ph-IAA and 0.5 µM AGB1 for 24 h. The scale bars indicate 10 µm.

Interestingly, we found that the simultaneous depletion of ORC1 and CDC6 led to premature mitosis without DNA replication (**Fig. 7C, D**). In the rounded cells, histone H3 Ser10 was highly phosphorylated, showing they are in mitosis (**Fig. 7C**). In these cells, condensed chromosomes did not show sister chromatids; instead, they showed single-chromatids (**Fig. 7D**). Furthermore, these single-chromatid chromosomes failed to align at the metaphase plate, even though spindle formation normally occurred (**Fig. S7B**). This failure was likely due to the inability to establish a centromeric bipolar attachment to microtubules. We concluded that ORC1 and CDC6 co-depletion by the AID2-BromoTag double-degron system completely suppresses DNA replication, leading to uncoupling DNA replication from the cell cycle control (**Fig. S7C**).

## Discussion

In this paper, we initially compared AID2, dTAG and BromoTag to learn their similarities and differences (**Figs 2** and **S2**). While 5-Ph-IAA, dTAGv-1 and AGB1 effectively depleted the GFP reporter in HCT116 and hTERT-RPE1, dTAG-13 was not as effective, especially in HCT116, as reported in other cell lines by others (Li *et al*, 2019; Olsen *et al*., 2022). This is likely due to a relatively low expression of the CRL4–CRBN components in HCT116 cells (**Fig. S1B**). The expression levels of the responsible E3 ligase likely affected depletion efficiency and kinetics because 5-Ph-IAA induced more highly profound reporter depletion in hTERT-RPE1 cells than in HCT116 cells (**Figs 2C, D** and **S2B, C**). Indeed, OsTIR1(F74G) in hTERT-RPE1 is overexpressed by random integration, while it is moderately expressed from the AAVS1 locus in HCT116. Additionally, we found that the reversible recovery after depletion with dTAG and BromoTag was poor compared with AID2 (**Figs 2D** and **S2D**), which is consistent with another report (Bondeson *et al*., 2022). We hypothesize that target degradation continued because the sub-stoichiometric activity of PROTACs dTAGv-1 and AGB1 bound to the CRL2–VHL E3 ligase can continue to be effective even after washing the compounds out from the culture medium. It will be important to test how generally applicable such features of these potent degron-based PROTAC degraders will be across other cell types/conditions and ultimately in vivo. One point that should be noted is that we focused on targeting nuclear proteins in this study. While AID2, dTAG and BromoTag are reported to degrade proteins in the cytoplasm (Bond *et al*., 2021; Nabet *et al*., 2020; Yesbolatova *et al*., 2020), the degradation efficiency and depletion kinetics in the cytoplasm might be different compared to the nucleus due to the localization and activity of the responsible E3 ligase.

Controlling the expression of two proteins is useful when studying their epistatic relationship and functional redundancy. We demonstrated that two proteins can be independently as well as simultaneously depleted by utilizing AID2 and BromoTag. As a proof-of-concept experiment, we depleted cohesin RAD21 and condensin SMC2 with AID2 and BromoTag, respectively (**Fig. 3**). As expected, RAD21 or SMC2 depletion caused defects in sister chromatid cohesion and in forming rod-like chromosomes, respectively (**Fig. 3C**) (Boteva *et al*., 2020; Sonoda *et al*., 2001; Takagi *et al*., 2018; Yesbolatova *et al*., 2020). Interestingly, the mitotic chromosomes depleted of both RAD21 and SMC2 were highly disorganized compared with those depleted of only SMC2. Our interpretation was that this chromosomal phenotype was an additive effect of sister-chromatid cohesion and chromosomal axis formation. It will be interesting to reveal what actually happens to the mitotic chromosomes in RAD21- and SMC2-depleted cells by Hi-C or super-resolution imaging approaches. We succeeded in developing a tandem-degron system with mAID-BromoTag to enhance target depletion and accelerate depletion kinetics (**Figs 4, 5** and **S5**). Empirically, we have achieved a near-knockout phenotype with AID2 for many replication proteins (Klein *et al*., 2021; Lim *et al*., 2023; Liu *et al*., 2023; Saito *et al*., 2022). However, we have found that there were some cases, such as ORC1 and CDC6, in which we could not deplete them sufficiently to study phenotypic defects. In these instances, the tandem-degron system proved to be advantageous. Moreover, each degron system shows different degradation efficiency depending on the protein of interest (POI) (Bondeson *et al*., 2022). Therefore, combining two degron systems is expected to effectively deplete more POIs than a single degron. We also found that tandem mAID tags (2mAID and 3mAID) were also effective for ORC1, causing strong defects in proliferation and MCM loading to chromatin DNA (**Fig. 6**). However, these tandem tags are intrinsically unstable and reduce the basal expression level of ORC1 independent of the presence or absence of OsTIR1(F74G) (**Figs 6A** and **S6A**). We previously reported that 3mAID was more effective than 2mAID in budding yeast (Kubota *et al*., 2013; Nishimura & Kanemaki, 2014). However, ORC1 was more effectively depleted with 2mAID than with 3mAID in human cells (**Fig. 6A**). Therefore, the tandem 2mAID tag can be used if the reduced expression level of a POI without 5-Ph-IAA does not cause a problem. By optimizing the degron and linker, it might be possible to create a more stable tandem mAID tag for AID2 in the future.

We demonstrated that MCM2-depleted cells by AID2 showed a lethal phenotype (**Fig. 1B, C**). On the other hand, we found that cells survived with a small amount of ORC1 and CDC6, and we needed to further deplete them by utilizing a mAID-BromoTag double degron to observe lethality (**Figs 1B, C, 5A, D** and **S5A, D**). These results clearly indicate that the level of depletion required to cause phenotypic defects is variable even though MCM2, ORC1 and CDC6 are involved in DNA replication. The ORC1–6 complex and CDC6 are all required for loading the MCM2– 7 complex to chromatin DNA (Costa & Diffley, 2022; Li *et al*, 2023), and a single ORC1–6 and CDC6 can potentially load multiple MCM double hexamers. Conversely, the loaded MCM double hexamers are converted to form the replicative CMG helicases for making the replication forks, and excess MCM2–7 is required for robust DNA replication (Ge *et al*, 2007; Ibarra *et al*, 2008; Peycheva *et al*, 2022). Their differential roles in DNA replication likely define the minimal amount needed for cell survival. In this study, we succeeded in effectively depleting ORC1 and CDC6 utilizing the double-degron system, causing defects in MCM loading and a lethal phenotype (**Figs 5** and **S5**). Interestingly, after ORC1 or CDC6 depletion, cells entered S phase with a small amount of MCM2–7 on chromatin (**Figs 5E** and **S5E**) and arrested in late-S and G2 phases, accumulating DNA damage (**Fig. S4**). Our interpretation is that HCT116 cells do not have the licensing checkpoint even though they are a p53-positive cell line, as we previously reported (McIntosh & Blow, 2012; Saito *et al*., 2022). These cells did not sense the lack of chromatin-bound MCM2–7 in a single cell cycle, entered S phase and then activated the DNA damage checkpoint. Because the double-degron system depletes the target very fast, it can be used for studying whether the licensing checkpoint exists in other cell lines. Regarding the conflicting reports on whether ORC1 is essential for DNA replication (Chou *et al*., 2021; Shibata *et al*., 2016), our data support that the conclusion that ORC1 is essential for loading MCM2–7 (**Figs 5** and **6**). Furthermore, we succeeded in demonstrating DNA replication was completely suppressed in cells depleted of ORC1 and CDC6 simultaneously, leading the cells to premature mitosis without DNA replication (**Fig. 7**). Therefore, we concluded DNA replication was artificially uncoupled from the cell cycle control (**Fig. S7C**), as shown by another report (Lemmens *et al*., 2018). This result suggests that there is no cellular system monitoring the completion of DNA replication. Instead, cells monitor the presence of replication forks via the DNA damage checkpoint pathway (Fragkos *et al*, 2015).

In summary, we described a comparison of AID2, dTAG and BromoTag, and our findings demonstrate that AID2 exhibited superior performance in terms of depletion efficiency, kinetics and reversible expression recovery in HCT116 and hTERT-RPE1 cells. Furthermore, we created new degron tools to regulate two proteins and enhance target depletion. To show the power of our new tools, we demonstrated that both ORC1 and CDC6 play an essential role in loading MCM2–7 and showed complete suppression of DNA replication. We anticipate that the new degron tools presented in this study will be useful in various fields of cell biology and contribute to future studies.

## Materials & Methods

### Plasmids

All plasmids used in this study and other new tagging plasmids are listed in **Table S1**. These plasmids and their sequence information are available from Addgene. ***Cell lines***

All cell lines used in this study and their genotype are listed in **Table S2**.

### Cell culture

HCT116 cells were cultured in McCoy’s 5A, supplemented with 10% FBS, 2 mM L-glutamine, 100 U/ml penicillin and 100 µg/mL streptomycin at 37 °C in a 5% CO_2_ incubator. To construct the GFP reporter cells expressing dTAG, BromoTag and mAID degrons tandemly, parental HCT116 cells expressing OsTIR1(F74G) were co-transfected with pMK459 (dTAG-BromoTag-mAID-EGFP-NLS) and pCMV-hyBase (PiggyBac transposase) using ViaFect Transfection reagent (Promega, #E498A) and Opti-MEM reduced serum medium (Thermo Fisher Scientific, #31985062) in a 12-well plate following the manufacturer’s instruction (Yusa *et al*, 2011). Cells were selected with 100 µg/mL Hygromycin B Gold (Invivogen, #ant-hg-5). Selected clones were isolated and GFP expression was confirmed by Western blotting and flow cytometry.

hTERT RPE1 cells were cultured in D-MEM/Ham’s F-12 with L-glutamine and Phenol Red, supplemented with 10% FBS, 100 U/ml penicillin and 100 µg/mL streptomycin at 37 °C in a 5% CO_2_ incubator. Initially, we established an hTERT RPE1 stable cell line expressing OsTIR1(F74G) by introducing pMK444 (EF1-OsTIR1(F74G)) and pCMV-hyBase (Yusa *et al*., 2011). hTERT RPE1 cells expressing OsTIR1(F74G) were co-transfected with pMK467 (dTAG-BromoTag-mAID-EGFP-NLS) and pCS-TP (TOL2 transposase) with the Neon system (Thermo Fisher Scientific) (Sumiyama *et al*, 2010). Cells were selected with 100 μg/mL Hygromycin B Gold. Selected clones were isolated and GFP expression was confirmed by Western blotting and flow cytometry.

All HCT116 cell lines expressing degron-fused protein(s) were generated following the protocol previously published (Saito & Kanemaki, 2021). Briefly, we constructed donor and CRISPR plasmids for tagging each gene. These CRISPR plasmids were designed to target MCM2 (5’-GAGGCCCTATGCCATCC/ATA-3’), ORC1 (5’-CCTTGTGGGGTAGTGTG/CCA-3’), CDC6 (5’-CATGCCTCAAACCCGAT/CCC-3’), RAD21 (5’-CCAAGGTTCCATATTAT/ATA-3’) and SMC2 (5’-ACCACCCAAAGGAGCAC/ATG-3’). After transecting both plasmids, cells were selected in the presence of an appropriate antibiotic. Single colonies were isolated and clones containing bi-allelic insertion at the target gene loci were selected by genomic PCR. The expression of degron-fused target protein was confirmed by Western blotting.

### Degrader compounds

5-Ph-IAA was synthesized as previously described (Yesbolatova *et al*., 2020). AGB1 for the initial studies was synthesized as previously reported (Bond *et al*., 2021), and later commercially obtained (Tocris #7686). dTAG-13 and dTAGv-1 were commercially obtained (Tocris #6605 and 6914, respectively).

### Colony formation assay

In a 6-well plate, 1500 cells were seeded and cultured in the presence or absence of indicated ligands for 7 days. The culture medium was exchanged for fresh medium containing appropriate ligands on day 4. Cells were fixed and stained with crystal violet solution (6.0% Glutaraldehyde, 0.5% crystal violet).

### Flow cytometric analysis

Reporter cells in the HCT116 or hTERT RPE1 background were seeded in a six-well plate, except for HCT116 RAD21-mACl/SMC2-BTCh cells which were seeded in a 12-well plate. They were grown for 2 days to obtain 80–90% confluency and treated with ligands as indicated in the figure legends. For detecting GFP/mClover and mCherry signals after ligand treatment, cells were trypsinized and fixed in 4% methanol-free paraformaldehyde (PFA) at 4 °C for 20 min or overnight. Fixed cells were washed and resuspended in 1% BSA/PBS. Flow cytometric analysis was performed on a BD Accuri C6 (BD Biosciences) or a BD FACSCelesta (BD Biosciences). Ten thousand cells were analysed from each sample with FlowJo 10.9.0 software (BD Biosciences).

To monitor chromatin-bound MCM2 (**Figs 5, 6** and **S5**), cells were permeabilized with 0.2% Triton X-100 in PBS on ice for 2 min and washed with ice-cold PBS twice. The pre-extracted cells were fixed with 4% PFA for 20 min at 4 °C. The fixed cells were washed with BSA/PBS-T (1% BSA and 0.2% Tween 20 in PBS) and blocked with 5% normal goat serum in BSA/PBS-T for 30 min at RT. Subsequently, the cells were incubated with anti-MCM2 rabbit monoclonal antibody for 1.5 h at RT. After washing three times with BSA/PBS-T, the cells were incubated with anti-rabbit goat Alexa 647 antibody for 1 h at RT. Subsequently, the cells were washed three times with BSA/PBS-T and were incubated with 24 µg/mL of propidium iodide and 50 µg/mL of RNase A in 1% BSA/PBS for 30 min at RT. Fluorescent signals in the immuno-stained cells were detected by a BD Accuri C6 flow cytometer. We analysed 10,000 gated single cells for each sample with FlowJo 10.9.0 software. The background level of loaded MCM2 (black dots in **Figs 5E, 6C, 6D** and **S5E**) was determined by using a control sample without MCM2 staining.

### Immunoblotting

Cells were harvested by trypsin and washed with culture medium and PBS. Cells were lysed with RIPA buffer (25 mM Tris-HCl pH 7.5, 150 mM NaCl, 1% NP-40, 1% sodium deoxycholate, 0.1% SDS) containing complete protease inhibitor cocktail (Roche, #1187580001) for 30 min on ice. Subsequently, the tubes were centrifuged for 15 min at 4 °C and then the supernatant was mixed with the same amount of SDS-sample buffer (Cosmo Bio, #423420) before heating at 95 °C for 5 min. The denatured protein samples were separated on a 7.5% or 10% TGX Stain-Free gel (BioRad) and transferred to a nitrocellulose membrane (Cytiva, #10600003). The membrane was processed with 1% skim milk in TBS-T for 15 min and incubated with primary antibody at 4 °C overnight. After washing with TBS-T, the membrane was incubated with corresponding secondary antibody for 1 h at RT. Proteins were detected by ChemiDoc Touch MP imaging system (BioRad), and the detected signals were quantified using Image Lab software (ver. 6.0.1, BioRad).

### Antibodies

All antibodies used in this study are listed in **Table S3**.

### Chromosome spreads

RAD21-mACl/SMC2-BTCh cells were seeded in a six-well plate. Two to three days later, cells were cultured in the presence and absence of 1 µM 5-Ph-IAA and/or 0.5 µM AGB1 for 2.5 h. Subsequently, KaryoMAX Colcemid Solution (Thermo Fisher Scientific, #15212012) was added to the medium at a final concentration of 0.1 µg/mL and then cells were incubated for an additional 0.5 h. Cells were harvested, washed with culture medium and resuspended in 1 mL of pre-warmed 75 mM KCl before incubation at 37 °C for 15 min. Subsequently, 30 µL of MeOH/acetic acid (3:1) was added to the tube, and incubated for 5 min at RT. After centrifugation, the supernatant was removed, and the cell pellet was resuspended with 500 µL of MeOH/AA and incubated for 5 min at RT. This incubation step was repeated with fresh MeOH/AA. After centrifugation, cells were resuspended in 150 µL MeOH/AA and incubated at –30 °C until observation. Five microlitres of the resuspended sample was dropped onto a tilted slide grass and dried for up to 30 min at 60 °C. Dried nuclei were embedded with 5 µL of VECTASHIELD with DAPI (Vector Laboratories, #H-1200) in a coverslip.

### Microscopy

For immunofluorescence staining, cells were cultured on coverslips in the presence or absence of 1 µM 5-Ph-IAA and/or 0.5 µM AGB1 for the indicated time and fixed with 4% PFA for 20 min at RT. After washing with PBS-T, cells were permeabilized with 0.3% TritonX-100 in PBS for 5 min at RT. After washing, the cells were blocked with 5% BSA in PBS for 20 min at RT and incubated with an appropriate primary antibody for 1.5 h at RT. After washing, the cells were incubated with a secondary antibody and Hoechst 33342 for 1 h at RT. The coverslip with stained cells was mounted with 5 µL of ProLong Glass antifade mount (Thermo Fisher Scientific) on a slide glass. For detecting DNA replication by EdU incorporation, cells were cultured with 10 µM EdU for 30 min before fixation. Incorporated EdU was stained using the Click-iT EdU Imaging Kit (Thermo Fisher Scientific, #C10339) according to the manufacturer’s instruction. The fluorescent signals were captured with a Delta Vision Personal DV system (GE Healthcare) with an inverted microscope (IX71, Olympus) through a PlanApo 60×/1.42 oil immersion objective lens (Olympus). All pictures were deconvoluted. At least, 250 nuclei were processed for quantification of 53BP1 foci by using Volocity software (ver. 6.3.1, PerkinElmer). For observing mitotic spindles shown in Fig. S7B, images were taken by a FLUOVIEW FV3000 confocal microscope (Olympus) with a UPlanXApo 60x/1.42 oil immersion objective lens (Olympus).

## Acknowledgements

We thank all members of the Kanemaki laboratory for discussion and support. We also thank Ms Karina Polkovnychenko, Dr Yasukazu Daigaku and Dr Conner Craigon for the initial study as a NIG intern, for sharing the hTERT-RPE1 cell line expressing OsTIR1(F74G) and for providing a plasmid encoding BromoTag, respectively. We appreciate Dr Kazuhiro Maeshima and Ms Shiori Iida for the discussion on chromosome architecture. YH is a JSPS Research Fellow for Young Scientists (DC2), and MI is a MEXT scholarship fellow. This work was supported by JSPS KAKENHI (JP21H0419 and JP23H04925) and JST CREST (JPMJCR21E6) to MTK.

## Competing interests

AC is a scientific founder, shareholder and advisor of Amphista Therapeutics, a company that is developing targeted protein degradation therapeutic platforms. The Ciulli laboratory receives or has received sponsored research support from Almirall, Amgen, Amphista Therapeutics, Boehringer Ingelheim, Eisai, Merck KaaG, Nurix Therapeutics, Ono Pharmaceutical and Tocris-Biotechne. The other authors do not have any competing interests.

## References

Bekker-Jensen DB, Kelstrup CD, Batth TS, Larsen SC, Haldrup C, Bramsen JB, Sorensen KD, Hoyer S, Orntoft TF, Andersen CL et al (2017) An Optimized Shotgun Strategy for the Rapid Generation of Comprehensive Human Proteomes. Cell Syst 4: 587–599 e584

Bond AG, Craigon C, Chan KH, Testa A, Karapetsas A, Fasimoye R, Macartney T, Blow JJ, Alessi DR, Ciulli A (2021) Development of BromoTag: A “Bump-and-Hole”-PROTAC System to Induce Potent, Rapid, and Selective Degradation of Tagged Target Proteins. J Med Chem 64: 15477–15502

Bondeson DP, Mullin-Bernstein Z, Oliver S, Skipper TA, Atack TC, Bick N, Ching M, Guirguis AA, Kwon J, Langan C et al (2022) Systematic profiling of conditional degron tag technologies for target validation studies. Nat Commun 13: 5495

Boteva L, Nozawa RS, Naughton C, Samejima K, Earnshaw WC, Gilbert N (2020) Common Fragile Sites Are Characterized by Faulty Condensin Loading after Replication Stress. Cell Rep 32: 108177

Bouguenina H, Nicolaou S, Le Bihan YV, Bowling EA, Calderon C, Caldwell JJ, Harrington B, Hayes A, McAndrew PC, Mitsopoulos C et al (2023) iTAG an optimized IMiD-induced degron for targeted protein degradation in human and murine cells. iScience 26: 107059

Buckley DL, Raina K, Darricarrere N, Hines J, Gustafson JL, Smith IE, Miah AH, Harling JD, Crews CM (2015) HaloPROTACS: Use of Small Molecule PROTACs to Induce Degradation of HaloTag Fusion Proteins. ACS chemical biology 10: 1831–1837

Chou HC, Bhalla K, Demerdesh OE, Klingbeil O, Hanington K, Aganezov S, Andrews P, Alsudani H, Chang K, Vakoc CR et al (2021) The human origin recognition complex is essential for pre-RC assembly, mitosis, and maintenance of nuclear structure. Elife 10

Costa A, Diffley JFX (2022) The Initiation of Eukaryotic DNA Replication. Annu Rev Biochem 91: 107–131

Elbashir SM, Harborth J, Lendeckel W, Yalcin A, Weber K, Tuschl T (2001) Duplexes of 21-nucleotide RNAs mediate RNA interference in cultured mammalian cells. Nature 411: 494–498

Evrin C, Alvarez V, Ainsworth J, Fujisawa R, Alabert C, Labib KP (2023) DONSON is required for CMG helicase assembly in the mammalian cell cycle. EMBO Rep 24: e57677

Fragkos M, Ganier O, Coulombe P, Mechali M (2015) DNA replication origin activation in space and time. Nature reviews Molecular cell biology 16: 360–374

Ge XQ, Jackson DA, Blow JJ (2007) Dormant origins licensed by excess Mcm2-7 are required for human cells to survive replicative stress. Genes & development 21: 3331–3341

Gu H, Zou YR, Rajewsky K (1993) Independent control of immunoglobulin switch recombination at individual switch regions evidenced through Cre-loxP-mediated gene targeting. Cell 73: 1155–1164

Haland TW, Boye E, Stokke T, Grallert B, Syljuasen RG (2015) Simultaneous measurement of passage through the restriction point and MCM loading in single cells. Nucleic Acids Res 43: e150

Ibarra A, Schwob E, Mendez J (2008) Excess MCM proteins protect human cells from replicative stress by licensing backup origins of replication. Proceedings of the National Academy of Sciences of the United States of America 105: 8956–8961

Jaeger MG, Winter GE (2021) Fast-acting chemical tools to delineate causality in transcriptional control. Molecular cell 81: 1617–1630

Jeppsson K, Kanno T, Shirahige K, Sjogren C (2014) The maintenance of chromosome structure: positioning and functioning of SMC complexes. Nature reviews Molecular cell biology 15: 601–614

Kanemaki MT (2022) Ligand-induced degrons for studying nuclear functions. Curr Opin Cell Biol 74: 29–36

Kleiger G, Mayor T (2014) Perilous journey: a tour of the ubiquitin-proteasome system. Trends Cell Biol 24: 352–359

Klein KN, Zhao PA, Lyu X, Sasaki T, Bartlett DA, Singh AM, Tasan I, Zhang M, Watts LP, Hiraga SI et al (2021) Replication timing maintains the global epigenetic state in human cells. Science 372: 371–378

Klemm RD, Bell SP (2001) ATP bound to the origin recognition complex is important for preRC formation. Proceedings of the National Academy of Sciences of the United States of America 98: 8361–8367

Koduri V, McBrayer SK, Liberzon E, Wang AC, Briggs KJ, Cho H, Kaelin WG, Jr. (2019) Peptidic degron for IMiD-induced degradation of heterologous proteins. Proceedings of the National Academy of Sciences of the United States of America 116: 2539–2544

Kubota T, Nishimura K, Kanemaki MT, Donaldson AD (2013) The Elg1 replication factor C-like complex functions in PCNA unloading during DNA replication. Molecular cell 50: 273–280

Lemmens B, Hegarat N, Akopyan K, Sala-Gaston J, Bartek J, Hochegger H, Lindqvist A (2018) DNA Replication Determines Timing of Mitosis by Restricting CDK1 and PLK1 Activation. Molecular cell 71: 117–128 e113

Li J, Dong J, Wang W, Yu D, Fan X, Hui YC, Lee CSK, Lam WH, Alary N, Yang Y et al (2023) The human pre-replication complex is an open complex. Cell 186: 98–111 e121

Li S, Prasanna X, Salo VT, Vattulainen I, Ikonen E (2019) An efficient auxin-inducible degron system with low basal degradation in human cells. Nature methods 16: 866–869

Lim Y, Tamayo-Orrego L, Schmid E, Tarnauskaite Z, Kochenova OV, Gruar R, Muramatsu S, Lynch L, Schlie AV, Carroll PL et al (2023) In silico protein interaction screening uncovers DONSON’s role in replication initiation. Science 381: eadi3448

Liu W, Saito Y, Jackson J, Bhowmick R, Kanemaki MT, Vindigni A, Cortez D (2023) RAD51 bypasses the CMG helicase to promote replication fork reversal. Science 380: 382–387

McIntosh D, Blow JJ (2012) Dormant origins, the licensing checkpoint, and the response to replicative stresses. Cold Spring Harb Perspect Biol 4

Morawska M, Ulrich HD (2013) An expanded tool kit for the auxin-inducible degron system in budding yeast. Yeast 30: 341–351

Mylonas C, Lee C, Auld AL, Cisse, II, Boyer LA (2021) A dual role for H2A.Z.1 in modulating the dynamics of RNA polymerase II initiation and elongation. Nature structural & molecular biology 28: 435–442

Nabet B, Ferguson FM, Seong BKA, Kuljanin M, Leggett AL, Mohardt ML, Robichaud A, Conway AS, Buckley DL, Mancias JD et al (2020) Rapid and direct control of target protein levels with VHL-recruiting dTAG molecules. Nat Commun 11: 4687

Nabet B, Roberts JM, Buckley DL, Paulk J, Dastjerdi S, Yang A, Leggett AL, Erb MA, Lawlor MA, Souza A et al (2018) The dTAG system for immediate and target-specific protein degradation. Nature chemical biology 14: 431–441

Natsume T, Kiyomitsu T, Saga Y, Kanemaki MT (2016) Rapid Protein Depletion in Human Cells by Auxin-Inducible Degron Tagging with Short Homology Donors. Cell Rep 15: 210–218

Nishimura K, Fukagawa T, Takisawa H, Kakimoto T, Kanemaki M (2009) An auxin-based degron system for the rapid depletion of proteins in nonplant cells. Nature methods 6: 917–922

Nishimura K, Kanemaki MT (2014) Rapid Depletion of Budding Yeast Proteins via the Fusion of an Auxin-Inducible Degron (AID). Curr Protoc Cell Biol 64: 20 29 21–16

Nishimura K, Yamada R, Hagihara S, Iwasaki R, Uchida N, Kamura T, Takahashi K, Torii KU, Fukagawa T (2020) A super-sensitive auxin-inducible degron system with an engineered auxin-TIR1 pair. Nucleic Acids Res 48: e108

Noviello G, Gjaltema RAF, Schulz EG (2023) CasTuner is a degron and CRISPR/Cas-based toolkit for analog tuning of endogenous gene expression. Nat Commun 14: 3225

Nowak RP, Xiong Y, Kirmani N, Kalabathula J, Donovan KA, Eleuteri NA, Yuan JC, Fischer ES (2021) Structure-Guided Design of a “Bump-and-Hole” Bromodomain-Based Degradation Tag. J Med Chem 64: 11637–11650

Olsen SN, Godfrey L, Healy JP, Choi YA, Kai Y, Hatton C, Perner F, Haarer EL, Nabet B, Yuan GC et al (2022) MLL::AF9 degradation induces rapid changes in transcriptional elongation and subsequent loss of an active chromatin landscape. Molecular cell 82: 1140–1155 e1111

Peycheva M, Neumann T, Malzl D, Nazarova M, Schoeberl UE, Pavri R (2022) DNA replication timing directly regulates the frequency of oncogenic chromosomal translocations. Science 377: eabj5502

Saito Y, Kanemaki MT (2021) Targeted Protein Depletion Using the Auxin-Inducible Degron 2 (AID2) System. Curr Protoc 1: e219

Saito Y, Santosa V, Ishiguro KI, Kanemaki MT (2022) MCMBP promotes the assembly of the MCM2-7 hetero-hexamer to ensure robust DNA replication in human cells. Elife 11

Shibata E, Kiran M, Shibata Y, Singh S, Kiran S, Dutta A (2016) Two subunits of human ORC are dispensable for DNA replication and proliferation. Elife 5

Sonoda E, Matsusaka T, Morrison C, Vagnarelli P, Hoshi O, Ushiki T, Nojima K, Fukagawa T, Waizenegger IC, Peters JM et al (2001) Scc1/Rad21/Mcd1 is required for sister chromatid cohesion and kinetochore function in vertebrate cells. Developmental cell 1: 759–770

Sumiyama K, Kawakami K, Yagita K (2010) A simple and highly efficient transgenesis method in mice with the Tol2 transposon system and cytoplasmic microinjection. Genomics 95: 306–311

Takagi M, Ono T, Natsume T, Sakamoto C, Nakao M, Saitoh N, Kanemaki MT, Hirano T, Imamoto N (2018) Ki-67 and condensins support the integrity of mitotic chromosomes through distinct mechanisms. Journal of cell science 131

Weintraub AS, Li CH, Zamudio AV, Sigova AA, Hannett NM, Day DS, Abraham BJ, Cohen MA, Nabet B, Buckley DL et al (2017) YY1 Is a Structural Regulator of Enhancer-Promoter Loops. Cell 171: 1573–1588 e1528

Yamanaka S, Shoya Y, Matsuoka S, Nishida-Fukuda H, Shibata N, Sawasaki T (2020) An IMiD-induced SALL4 degron system for selective degradation of target proteins. Commun Biol 3: 515

Yesbolatova A, Saito Y, Kitamoto N, Makino-Itou H, Ajima R, Nakano R, Nakaoka H, Fukui K, Gamo K, Tominari Y et al (2020) The auxin-inducible degron 2 technology provides sharp degradation control in yeast, mammalian cells, and mice. Nat Commun 11: 5701

Yusa K, Zhou L, Li MA, Bradley A, Craig NL (2011) A hyperactive piggyBac transposase for mammalian applications. Proceedings of the National Academy of Sciences of the United States of America 108: 1531–1536

