## Supplementary Figures 1-7 for "Combination of AID2 and BromoTag expands the utility of degron-based protein knockdowns"

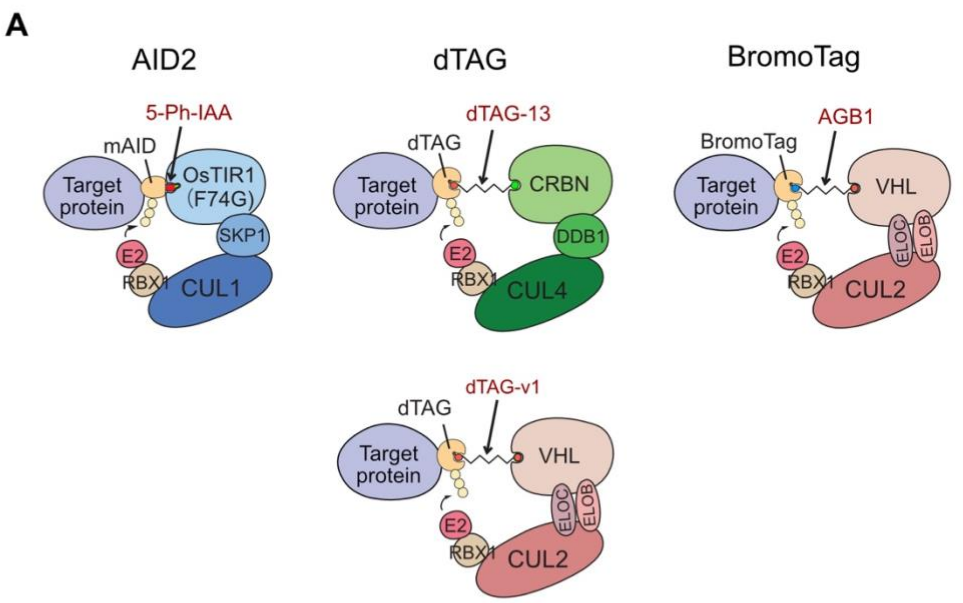


| Degron | mini-AID (mAID)<br>= IAA17(65–132aa) | dTAG<br>= FKBP12(F36V) | BromoTag<br>= BRD4-BD2(L387A) |
|---|---|---|---|
| Degron size | 7.5 kDa | 11.7 kDa | 14.8 kDa |
| Ligand | 5-Ph-IAA | dTAG-13, dTAG-v1 | AGB1 |
| E3 ligase | CRL1–OsTIR1(F74G) | CRL4–CRBN, CRL2–VHL | CRL2–VHL |

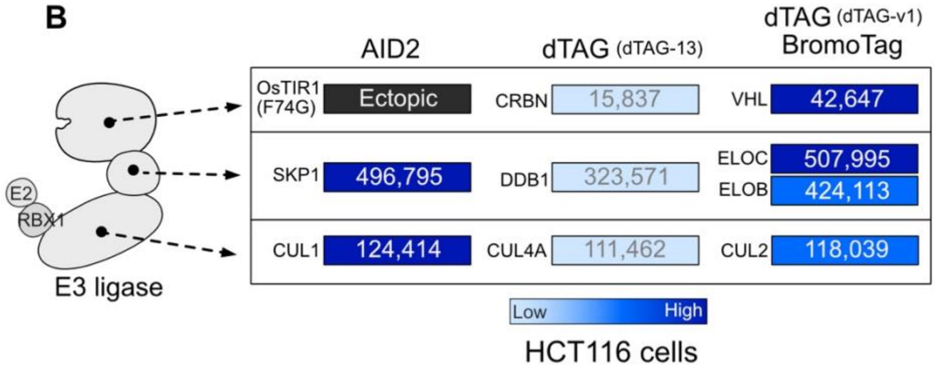


**Figure S1**

(**A**) Schematic illustration showing the AID2, dTAG and BromoTag systems. The table shows degron size, inducing ligand and the E3 ligase involved in target degradation. (**B**) The expression level of each E3 ligase component in HCT116 cells, originated from the paper by Bekker-Jensen et al {Bekker-Jensen, 2017 #323}.

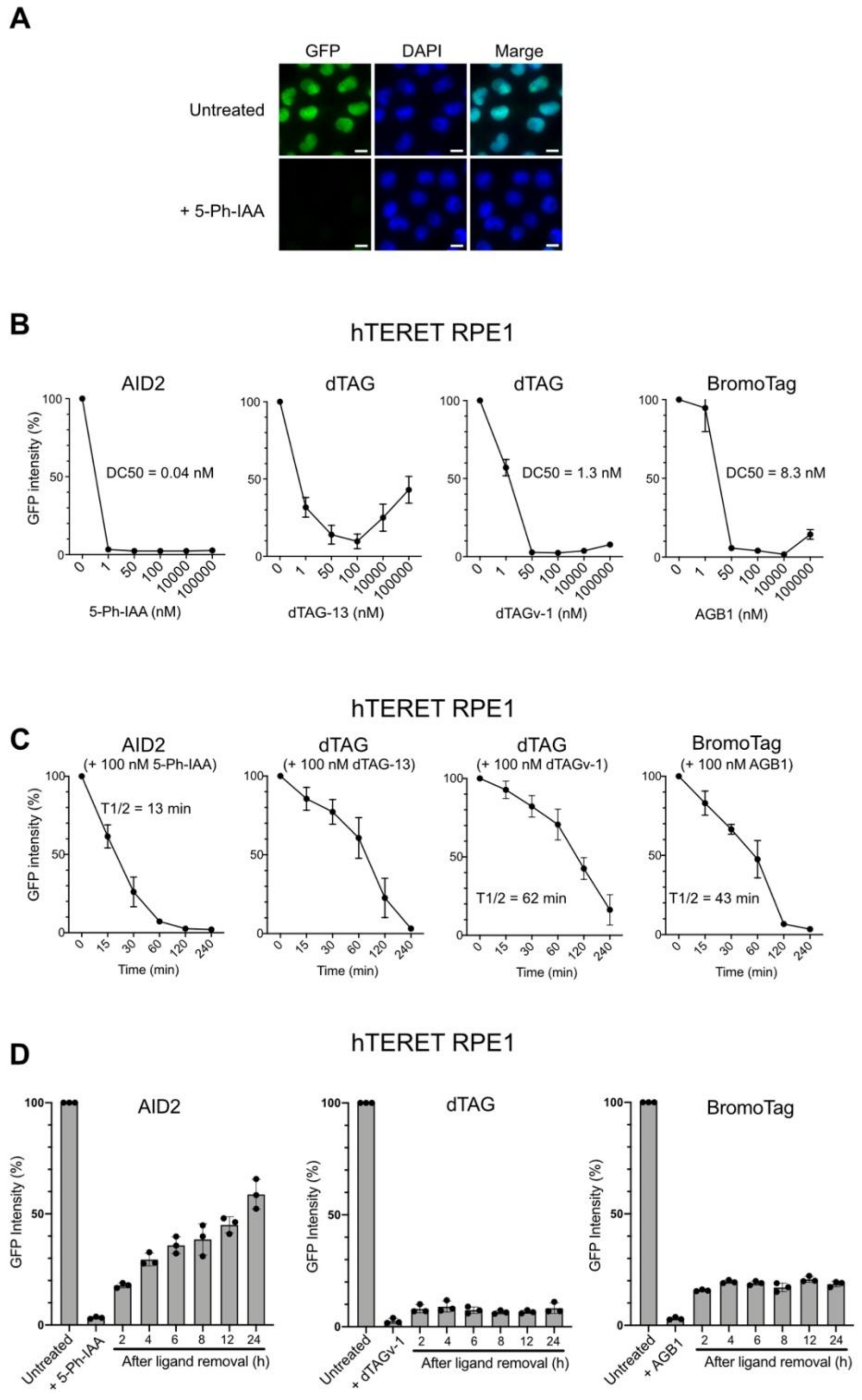


**Figure S2**

Comparison of the AID2, dTAG and BromoTag systems using a GFP reporter. (**A**) Representative fluorescent microscopic images of the HCT116 dTAG-BromoTag-mAID-EGFP-NLS reporter cells. The cells were treated with or without 1 µM 5-Ph-IAA for 2 h before fixation. The nuclei were stained with DAPI. Scale bars: 10 µm. (**B**) Dose-response of reporter depletion. The hTERT-RPE1 reporter cells were treated with the indicated concentrations of each ligand for 4 h. GFP intensity was analyzed taking the mock-treated cells as 100% (mean ± SD, n = 3). The DC50 values were calculated with the non-linear regression model on Graphpad Prism 8. (**C**) Time-course depletion of the reporter in hTERT-RPE1 cells. Cells were treated with 100 nM of the indicated ligand. Samples were taken at the indicated time point. GFP intensity was analyzed taking the mock-treated cells as 100% (mean ± SD, n = 3). The T1/2 was calculated with the non-linear regression model on Graphpad Prism 8. (**D**) Re-expression of the reporter after depletion by the AID2, dTAG or BromoTag system. The reporter cells were treated with 1nM 5-Ph-IAA, 100 nM dTAGv-1 or 100 nM AGB1 for 4 h before medium change. Samples were taken at the indicated time points, and the GFP intensity was analyzed by taking the mock-treated cells as 100% (mean± SD, n=3).

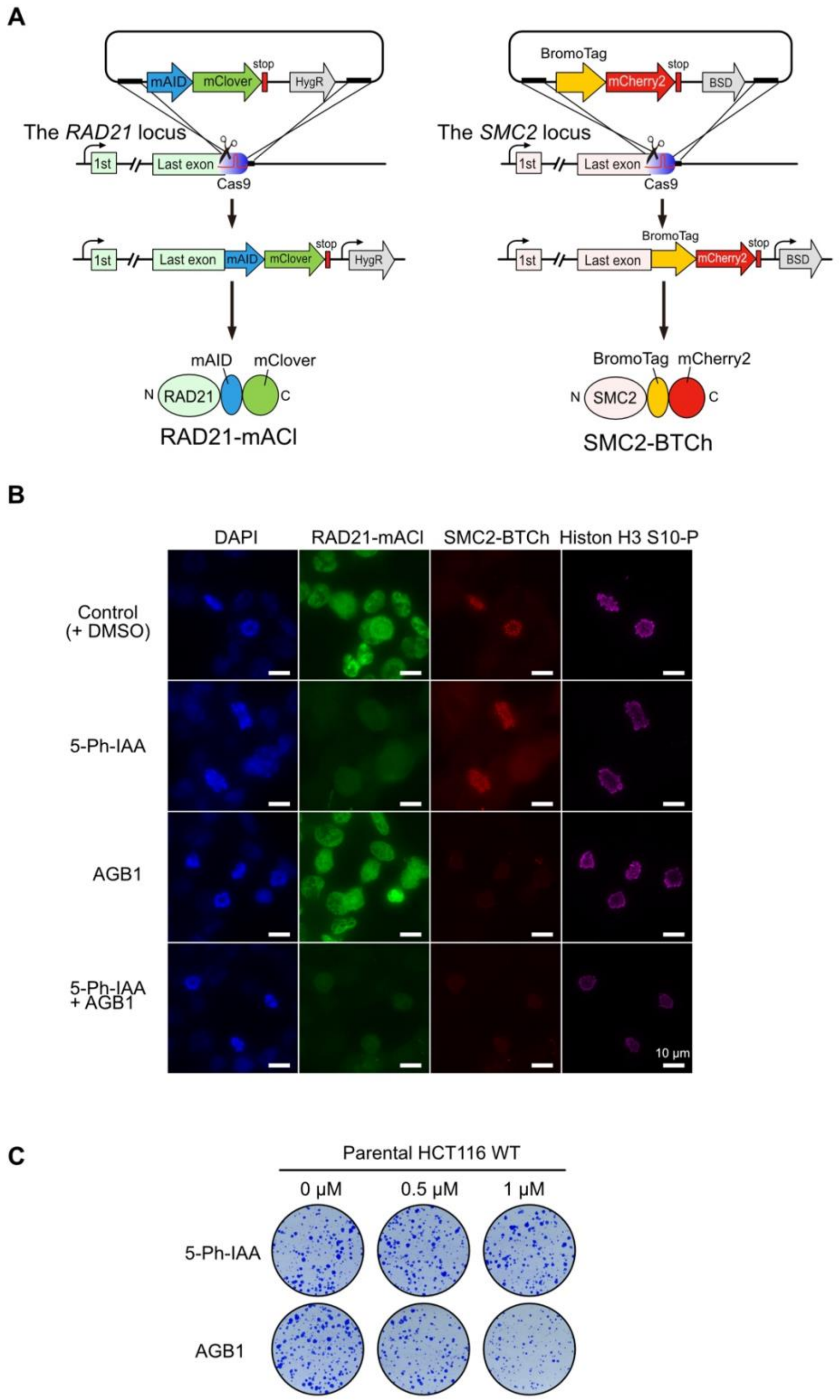


**Figure S3**

Related to Fig. 4. (**A**) Illustration showing the construction cells expressing RAD21-mAID-mClover (RAD21-mAC) and SMC2-BromoTag-mCherry2 (SMC2-BCh). (**B**) Fluorescent microscopic images of the RAD21-mAC/SMC2-BTCh cell line in the presence and absence of 1μM 5-Ph-IAA and/or 0.5 μM AGB1 for 4 h. Mitotic cells were stained with anti p-Histon H3 (Ser10) antibody. Ten slices taken every 0.5 µm were stacked. Scale bar: 10 µm. (**C**) Testing side effects of 5-Ph-IAA and AGB1 by colony formation. The parental HCT116 WT cells were cultured with the indicated dose of 5-Ph-IAA or AGB1 for 7 days. Colonies were stained with crystal violet.

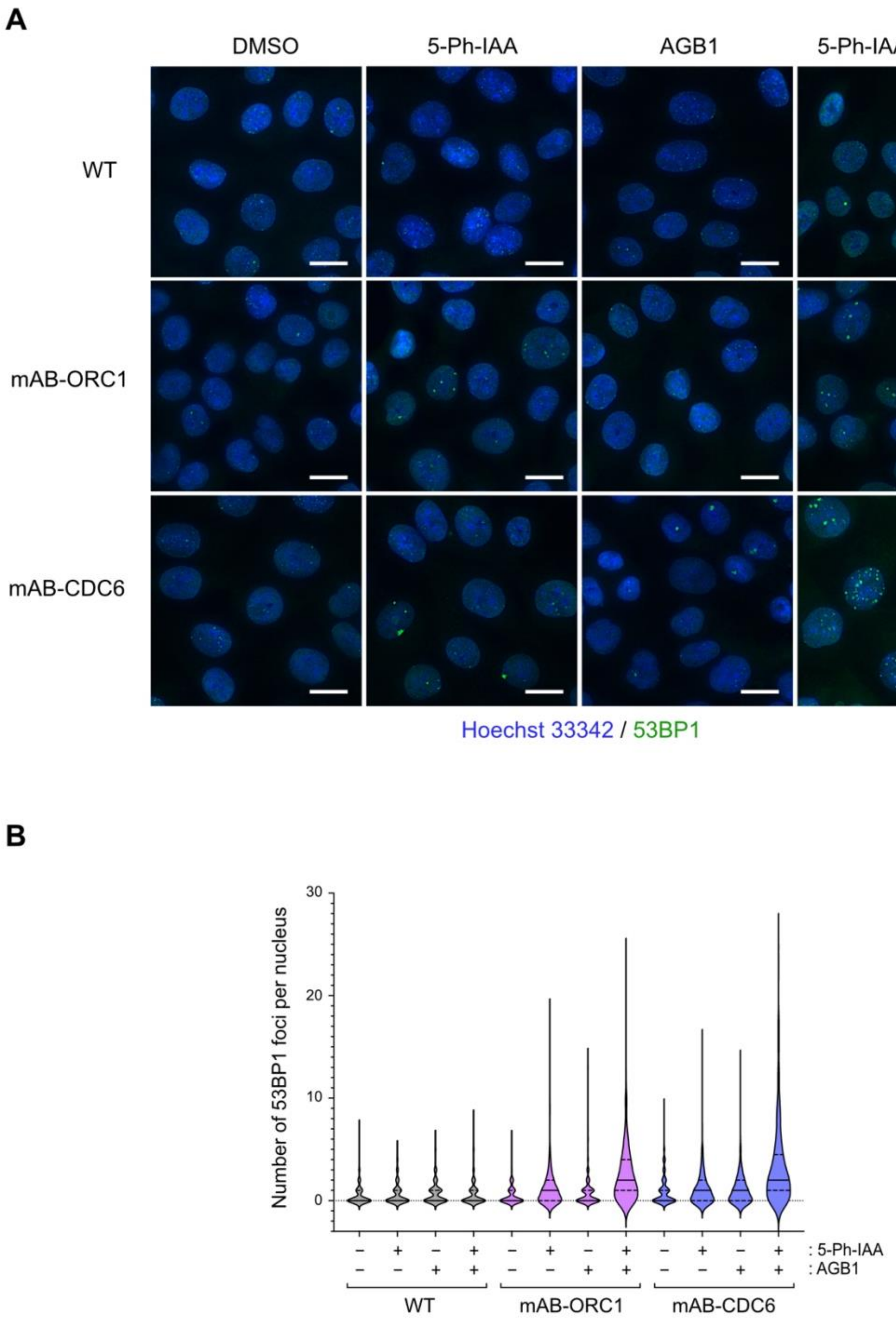


**Figure S4**

DNA damage foci formation in mAB-ORC1 and mAB-CDC6 cells. (**A**) Representative 53BP1 immunofluorescence images after treating the indicated cells with 1 µM 5-Ph-IAA, 0.5 µM AGB1 or both for 43 h. 53BP1 and DNA are shown in green and blue, respectively. (**B**) The number of 53BP1 foci per nucleus was quantified and presented in the violin plot. The solid lines show the median and dashed lines show the quartiles. More than 250 nuclei were quantified in each condition.

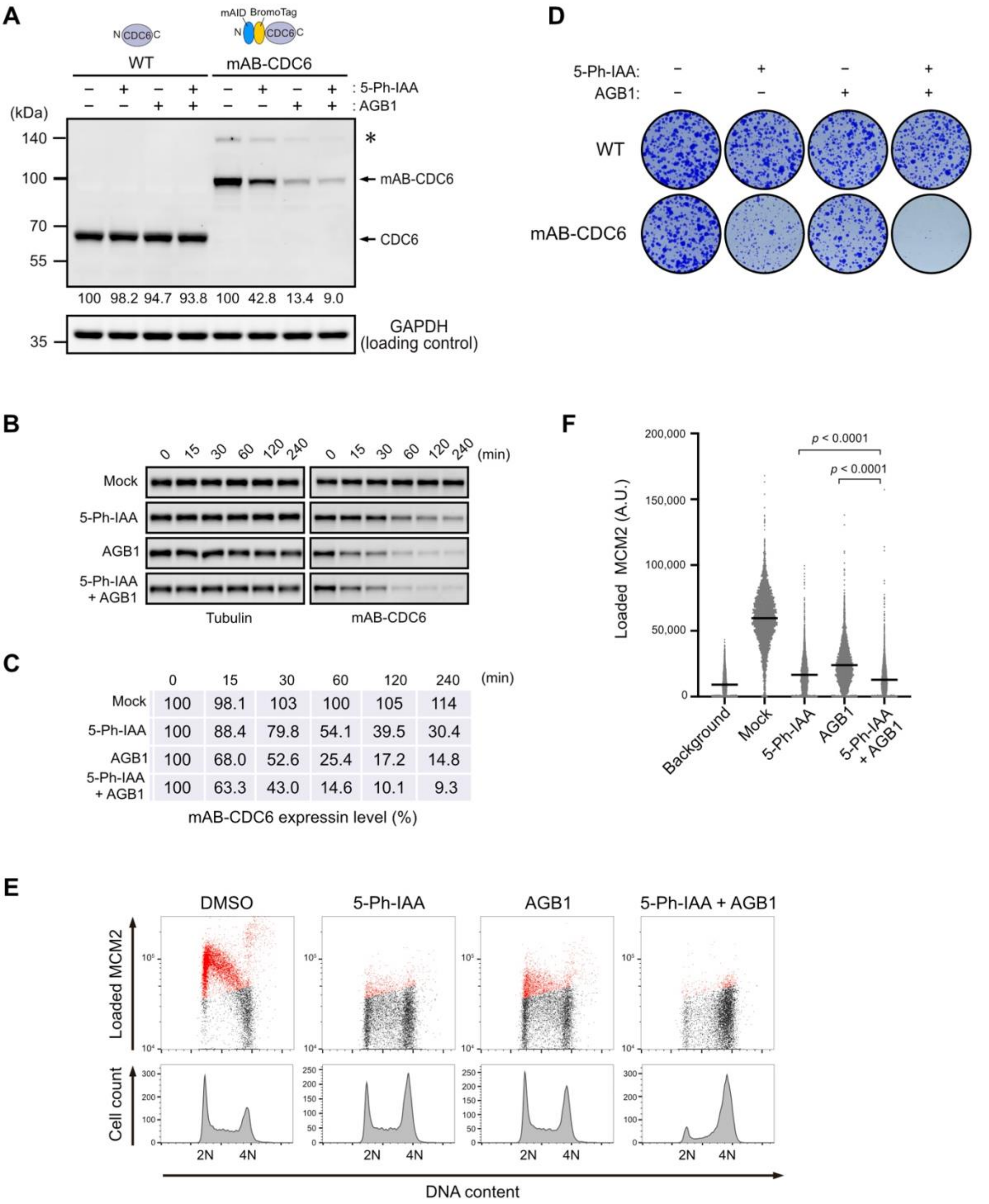


**Figure S5**

Double-degron with mAID and BromoTag enhances CDC6 depletion and confers profound defects in DNA replication. (**A**) The parental HCT116 wild-type (WT) and mAID-BromoTag-CDC6 (mAB-CDC6) cells were treated with 1 µM 5-Ph-IAA, 0.5 µM AGB1 or both for 2 h. Proteins were detected by anti-CDC6 and -GAPDH antibodies. Relative CDC6 levels taking the DMSO-treated control as 100% were shown under each blot. Each data was normalized with the corresponding tubulin loading control.

The asterisk indicates the HygR-P2A-mAID-BromoTag-CDC6 protein before self-cleavage at the P2A site. (**B**) Depletion kinetics of mAB-CDC6 in cells treated with 1 μM 5-Ph-IAA and/or 0.5 μM AGB1. Samples were taken at the indicated time points. (**C**) The blot data in panel B were quantified taking the 0 min sample as 100%. Each data was normalized with the corresponding tubulin loading control. (**D**) Colony formation of the parental HCT116 WT and mAB-CDC6 cells. Cells were cultured in the presence or absence of 1 μM 5-Ph-IAA and/or 0.5 μM AGB1 for 7 days. Colonies were stained with crystal violet. (**E**) (Upper panels) Levels of chromatin-loaded MCM2 and DNA in mAB-CDC6 treated with 1 μM 5-Ph-IAA and/or 0.5 μM AGB1 for 24 h. The MCM2-positive cells are shown in red. (Lower panels) Cell count histogram to the same samples. (**F**) Levels of chromatin-loaded MCM2 in mAB-CDC6 cells treated with the indicated ligand (bars: mean, n > 5,000 cells). Cells were synchronized in M phase with 50 ng/mL nocodazole for 14 h and released into a fresh media containing ligand. Cells were treated with 1 μM 5-Ph-IAA and/or 0.5 μM AGB1 2 h prior to nocodazole release. Samples were taken at 4 h after release when cells were in G1. Fixed cells stained without MCM2 antibody serve as the background. Statistical analysis was performed with Kruskal-Wllis test.

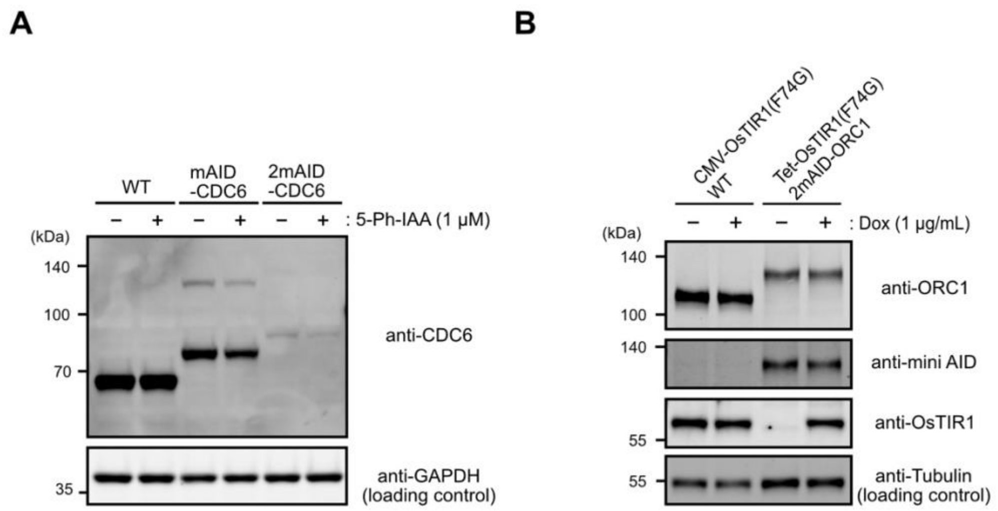


**Figure S6**

The 2mAID tag confers protein instability. (**A**) CDC6 level in the indicated cells treated with or without 1μM 5-Ph-IAA for 2 h. Proteins were detected with the indicated antibodies. (**B**) We generated a cell line expressing 2mAID-ORC1 in the Tet-OsTIR1(F74G) background, in which OsTIR1(F74G) is induced by the addition of doxycycline (Dox). The parental HCT116 WT cells used in this study (CMV-OsTIR1 background) and the 2mAID-ORC1 (Tet-OsTIR1(F74G) background) cells were treated with 1 μg/mL doxycycline for 24 h. Proteins were detected by the indicated antibodies.

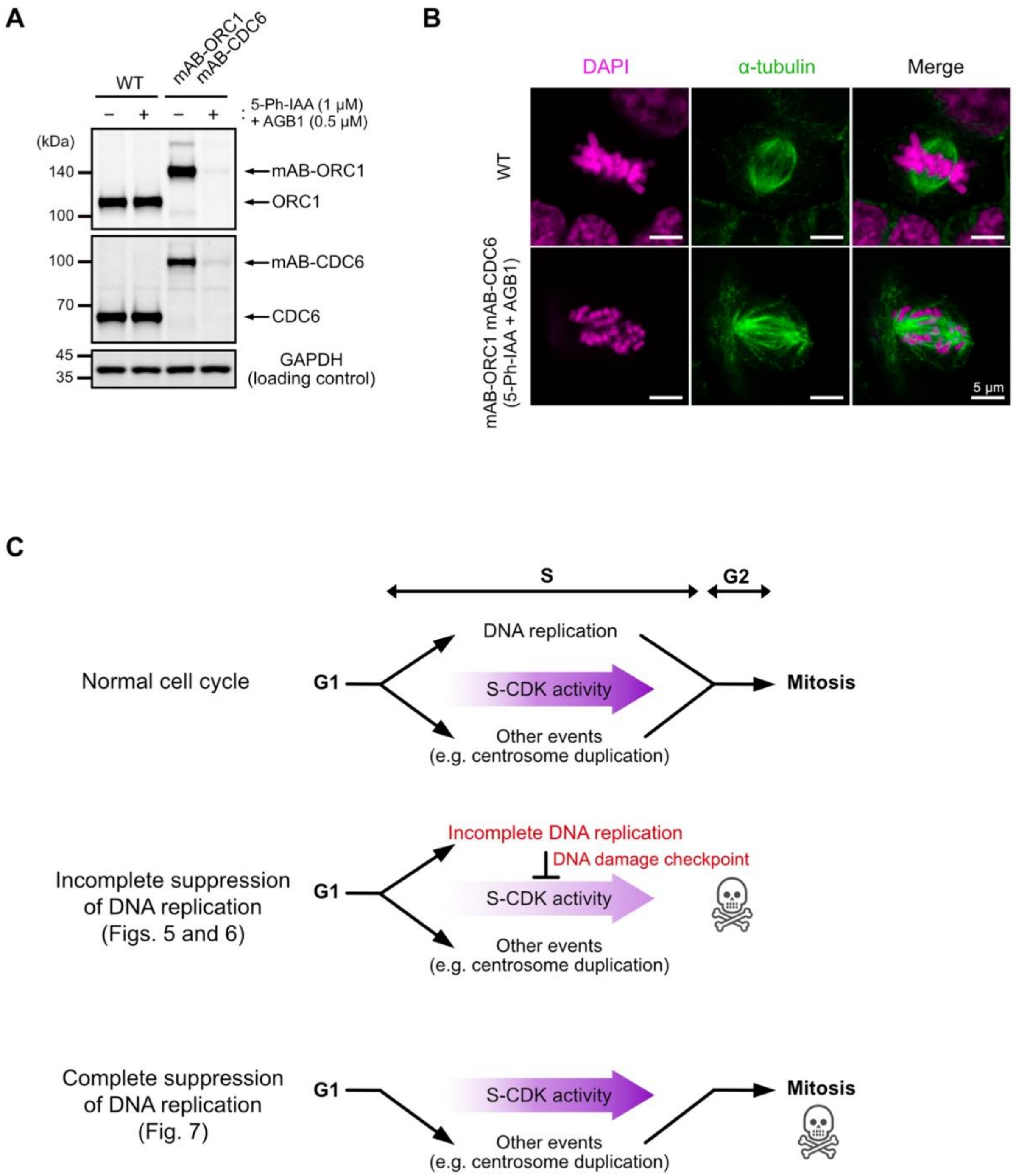


**Figure S7**

The mAB-ORC1 mAB-CDC6 cells enter mitosis without DNA replication after ORC1 and CDC6 co-depletion. (**A**) mAID-BromoTag was fused to the N-terminus of ORC1 and CDC6. The cells were treated with or without 1 μM 5-Ph-IAA and 0.5 μM AGB1 for 2 h before harvesting. Proteins were detected with anti-ORC1, -CDC6 and - GAPDH antibodies. (**B**) Confocal microscopic images of metaphase cells. Spindle and DNA were stained by anti-α-tubulin antibody and DAPI, respectively. HCT116 WT cells and the mAB-ORC1 mAB-CDC6 cells were cultured with 1 μM 5-Ph-IAA and 0.5 μM AGB1 for 24 h before fixation. (**C**) Illustration showing the relationship between DNA replication and the cell cycle control. Complete DNA suppression bypasses DNA replication resulting premature mitosis. Note that both incomplete and complete suppression of DNA replication leads to cell death. However, they were arrested at different cell cycle phases (late S/G2 and M, respectively).
